# Privacy-preserving search for chemical compound databases

**DOI:** 10.1101/013995

**Authors:** Kana Shimizu, Koji Nuida, Hiromi Arai, Shigeo Mitsunari, Nuttapong Attrapadung, Michiaki Hamada, Koji Tsuda, Takatsugu Hirokawa, Jun Sakuma, Goichiro Hanaoka, Kiyoshi Asai

## Abstract

**Background:** Searching for similar compounds in a database is the most important process for in-silico drug screening. Since a query compound is an important starting point for the new drug, a query holder, who is afraid of the query being monitored by the database server, usually downloads all the records in the database and uses them in a closed network. However, a serious dilemma arises when the database holder also wants to output no information except for the search results, and such a dilemma prevents the use of many important data resources.

**Results:** In order to overcome this dilemma, we developed a novel cryptographic protocol that enables database searching while keeping both the query holder’s privacy and database holder’s privacy. Generally, the application of cryptographic techniques to practical problems is difficult because versatile techniques are computationally expensive while computationally inexpensive techniques can perform only trivial computation tasks. In this study, our protocol is successfully built only from an additive-homomorphic cryptosystem, which allows only addition performed on encrypted values but is computationally efficient compared with versatile techniques such as general purpose multi-party computation. In an experiment searching ChEMBL, which consists of more than 1,200,000 compounds, the proposed method was 36,900 times faster in CPU time and 12,000 times as efficient in communication size compared with general purpose multi-party computation.

**Conclusion:** We proposed a novel privacy-preserving protocol for searching chemical compound databases. The proposed method, easily scaling for large-scale databases, may help to accelerate drug discovery research by making full use of unused but valuable data that includes sensitive information.

## Introduction

In recent years, the increasing cost of drug development and decreasing number of new chemical entities have become growing concerns [1]. One of the most popular approaches for overcoming these problems is searching for similar compounds in databases [2]. In order to improve the efficiency of this task, it is important to utilize as many data resources as possible. However, the following dilemma prevents the use of many existing data resources. Unpublished experimental results have been accumulated at many research sites, and such data has scientific value [3]. Since data holders are usually afraid of sensitive information leaking from the data resources, they do not want to release the full data, but they might allow authorized users to search the data as long as the users obtain only search results from which they cannot infer sensitive information. Likewise, private databases of industrial research might be made available if the sensitive information were sufficiently protected. On the other hand, query compounds are also sensitive information for the users, and thus the users usually avoid sending queries and want to download all of the data in order to conduct search tasks on their local computers. In short, we cannot utilize important data resources because both the data holder and the data user insist on their privacy. Therefore, an emerging issue is to develop novel technology that enables privacy-preserving similarity searches. We show several use cases in the next section.

Let us start by clarifying privacy problems in database searches. In a database search, two types of privacy are of concern: *“user privacy”* (also known as input privacy) and *“database privacy”* (also known as output privacy). The first is equal to protecting the user’s query from being leaked to others including the database holder. The second is equal to protecting the database contents from being leaked to others including the database user, except for the search results held by the user. Here we firstly consider the case of using no privacy-preserving techniques; namely, the user sends a plain query to the server and the server sends the search result. In this case, the user’s query is fully obtained by the server. On the database side, the server’s data is not directly leaked to the user. However, there is a potential risk that the user may infer the database contents from the search results. To protect user privacy, a scheme called single-database private information retrieval (PIR) has been proposed [4]. The simplest method for achieving PIR is that the user downloads all the contents of the database and searches on his/her local computer. Since this naive approach needs a huge communication size, several cryptographic techniques have been developed, in which the query is safely encrypted/randomized in the user’s computer and the database conducts the search without seeing the query. Although PIR is useful for searching public databases, it does not suit the purpose of searching private databases because of the lack of database privacy. Likewise, similarity evaluation protocols keep user privacy [5–7] but they do not sufficiently protect database privacy because the server directly outputs similarity scores that become important hints for inferring database contents.

Generally speaking, it is very difficult to keep both user privacy and database privacy, because the database side must prevent various attacks without seeing the user’s query. Among them, the following two attacks are major

- Regression attack Given one data point, the similarity between a target and the data point becomes a strong hint for detecting the target. The accuracy of the detection increases as the number of given data points becomes larger. In fact, a protocol that is not suitably designed may lead to even a small number of queries enabling the database user to detect the target. For example, when the server returns the exact distance between a query and a database entry, the range of the entry is rapidly narrowed as the number of queries increases, and the entry is finally detected uniquely by only almost the same number of queries as the dimension of the entry (see Fig. 1 for a detailed explanation). For example, in the case of using the MACCS keys, which are 166 bit structural key descriptors and often used for representing chemical compounds, a database entry is detected by sending only 166 queries. Therefore, it is necessary for the server to return the minimum information that is sufficient for the purpose of the search. In the Thresholding largely improves database privacy section, we will compare success probability of the regression attack for the case when the server returns minimum information (which our protocol aims for) and the other case when the server returns more information (which the previous method aims for).
- Illegal query attack Searching with an illegal query often causes unexpected server behaviour. In such a case, the server might return unexpected results that include important server information. To prevent this, the server should ensure the correctness of the user’s query.

**Figure 1:**
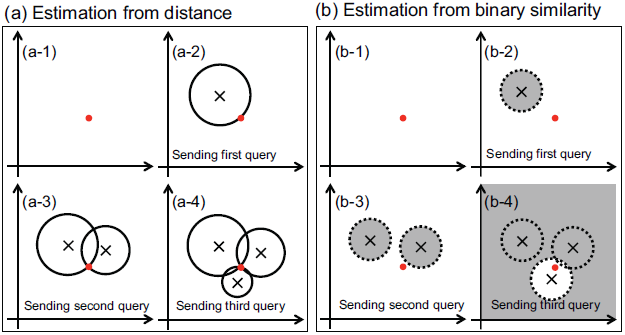
Schematic view showing a large difference in tolerance against the regression attack between two cases: (a) The server’s reply is the distance between the attacker’s query and the server’s data, (b) The server’s reply is the binary sign that shows whether or not the distance between the attacker’s query and the server’s data is larger than the given threshold. The red point represents the server’s data and x represents the attacker’s query. Prior to the query, the search spaces (white areas) in (a-1) and (b-1) are equal. After the first query has been sent, the search space in (a-2) is limited to the circle whose radius is the distance between the attacker’s query and the server’s data. On the other hand in (b-2), only the small area of the dashed circle whose radius is the given threshold (gray area) is excluded from the search space. By sending the second query, the attacker knows that one of the two intersections of the two circles in (a-3) is equal to the server’s data, while the search space is large in (b-3). Finally, the server’s data is detected by sending the third query in (a-4), however in (b-4), the search space is still large, even though the third query is within the given threshold.

A schematic view of the privacy-preserving database search problems discussed here is shown in Fig. 2.

**Figure 2:**
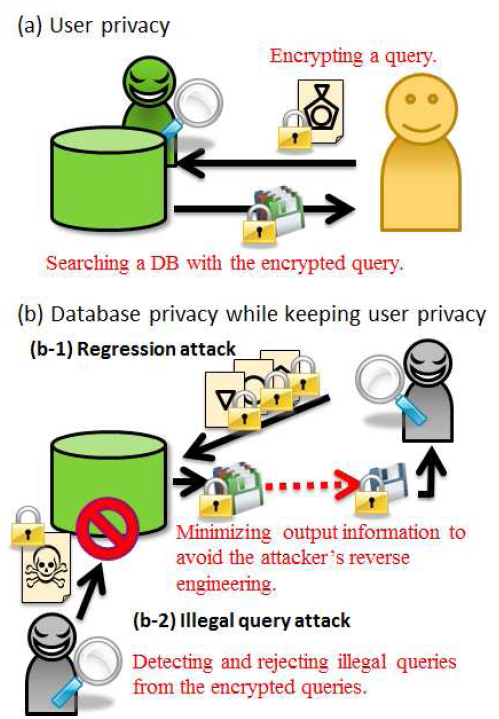
Schematic view of protection of (a) user privacy and (b) database privacy while keeping user privacy. For user privacy, the user’s query and the search result which includes the query information must be invisible to the database side during the search task. For database privacy, the server minimizes output information for preventing regression attacks (b-1), and also detects and rejects illegal queries that might cause unexpected information leakage (b-2). These server’s tasks must be carried out with the encrypted queries in order to keep user privacy.

In the field of cryptography, there have been studies of versatile techniques such as general purpose multi-party computation (GP-MPC) [8] and fully homomorphic encryption (FHE) [9], which enable the design of systems that maintain both user privacy and database privacy. However, these techniques require huge computational costs as well as intensive communications between the parties (see the recent performance evaluation of FHE [10]), so they are scarcely used in practical applications. In order to avoid using such techniques, a similarity search protocol using a trusted third party [11] and a privacy preserving SQL database using a trusted proxy server [12] have been proposed, but those methods assure privacy only when the third party does not collude with the user or the server, which is not convenient for many real problems. As far as we know, no practical method has been proposed despite the great importance of privacy-preserving similarity searching. To overcome this lack, we propose a novel privacy-preserving similarity search method that can strongly protect database privacy as well as user privacy while keeping a significantly low computational cost and small communication size.

The rest of this paper is organized as follows. In the next section, we summarize our achievements in this study. This is followed by the Cryptographic background section and the Method section, where we define the problem and introduce details of the proposed protocol. In the Security analyses section, both the user privacy and database privacy of the proposed protocol are discussed in detail. In the Performance evaluation section, the central processing unit (CPU) time and communication size of the proposed protocol are evaluated for two datasets extracted from ChEMBL. Finally, we present our conclusions for this study in the Conclusion section.

## Our Achievements

Here we focus on similarity search with the Tversky index of fingerprints, which is the most popular approach for chemical compound searches [13] and is used for various search problems in bioinformatics. To provide a concrete application, we address the problem of counting the number of similar compounds in a database, which solves various problems appearing in chemical compound searches. The following model describes the proposed method.

### Model 1.

*The user is a private chemical compound holder, and the server is a private database holder. The user learns nothing but the number of similar compounds in the server’s database, and the server learns nothing about the user’s query compound.*

Here we introduce only a small fraction of the many scientific or industrial problems solved by Model 1.

1. Secure pre-purchase inspection service for chemical compound. When a client considers the purchase of a commercial database such as a focused library [14], he/she wants to check whether the database includes a sufficient number of similar compounds, without sending his/her private query, but the server does not allow downloading of the database.
2. Secure patent compound filtering. When a client finds a new compound, he/she usually wants to know whether it infringes on competitors’ patents by searching the database of patent-protected compounds maintained by third parties. The same problem occurs when the client wants to check whether or not the compound is undesirable.
3. Secure negative results check. It is a common perception that current scientific publication is strongly biased against negative results [3], although a recent study showed statistically that negative results brought meaningful benefit [15]. Since researchers are reluctant to provide negative results, which often include sensitive information, a privacy-preserving system for sharing those results would greatly contribute to reducing redundant efforts for similar research topics. For example, it would be useful to have a system that allows a user to check whether the query is similar to failed compounds that have previously been examined in other laboratories.

In this study, we propose a novel protocol called the **secure similar compounds counter** (SSCC) which achieves Model 1. The first main achievement of this study is that SSCC is remarkably tolerant against regression attacks compared with existing protocols which directly output the similarity score. Moreover, we propose an efficient method for protecting the database from illegal query attacks. These points are discussed in the Security analyses section.

The second main achievement is that SSCC is significantly efficient both in computational cost and communication size. We carefully designed the protocol such that it uses only an additive-homomorphic cryptosystem, which is computationally efficient, and does not rely on any time-consuming cryptographic methods such as GP-MPC or FHE. Hence the performance of the protocol is sufficiently high for a large-scale database such as ChEMBL [16], as is shown in the Performance evaluation section.

## Cryptographic background

### Additively homomorphic encryption scheme

In this paper, we use an additive-homomorphic cryptosystem to design our protocol. The key feature of the additive-homomorphic cryptosystem is that it enables to perform additive operations on *encrypted* values. Therefore, intuitively, any standard computation algorithm can be converted into the privacy-preserving computation algorithm, if operations used in the standard algorithm can be replaced by additions.

More formally, we use a public-key encryption scheme (`Key Gen; Enc; Dec`), which is semantically secure; that is, an encryption result (ciphertext) leaks no information about the original message (plaintext) [17]. Here, `KeyGen` is a key generation algorithm for selecting a pair (`pk, sk`) of a public key `pk` and a secret key `sk`; `Enc`(*m*) denotes a ciphertext obtained by encrypting message *m* under the given `pk`; and `Dec`(*c*)denotes the decryption result of ciphertext *c* under the given `sk`. We also require the following additive-homomorphic properties:

- Given two ciphertexts `Enc`(*m*_1_) and `Enc`(*m*_2_) of messages *m*_1_ and *m*_2_, `Enc`(*m*_1_ + *m*_2_) can be computed without knowing *m*_1_, *m*_2_ and the secret key (denoted by `Enc`(*m*_1_) ⊕ `Enc`(*m*_2_)).
- Given a ciphertext `Enc`(*m*) of a message *m* and an integer *e*, `Enc`(*e* · *m*) can be computed without knowing *m* and the secret key(denoted by *e* ⊗ `Enc`(*m*)).

For example, we can use either the Paillier cryptosystem [18] or the “lifted” version of the ElGamal cryptosystem [19] as such an encryption scheme; now the second operation ⊗ can be achieved by repeating the first operation ⊕. We notice that the range of plaintexts for those cryptosystems can be naturally set as an integer interval [−*N*_1_*, N*_2_] for some sufficiently large *N*_1_*, N*_2_ *>* 0; therefore, the plaintexts are divided into positive ones, negative ones, and zero.

### Non-interactive zero-knowledge proof

Below, we discuss the following situation: A user (a prover) wants to make a server (a verifier) convinced that a ciphertext *c* generated by the user corresponds to a message *m* in {0, 1}, but does not want to reveal any information about which of 0 and 1 is *m*. This can be achieved by using a cryptographic tool called *non-interactive zero-knowledge* (*NIZK*) *proof*. In the present case, it enables the user to generate a “proof” associated with *c*, so that:

- If *m* is indeed in {0, 1}, then the server can verify this fact by testing the proof (without knowing *m* itself).
- If *m* ∉{0, 1}, then the user cannot generate a proof that passes the server’s test.
- The server cannot obtain any information about *m* from the proof, except for the fact that *m* ∈{0, 1}.

(See [20] for a general formulation.) Besides the existing general-purpose NIZK proofs, Sakai et al. [21] proposed an efficient scheme specific to the “lifted” ElGamal cryptosystem, which we use below. (See Section 1 of Additional File 1 in which we give the brief description of the NIZK proofs [21].)

## Method

The goal of this study is to design a protocol between a user and a server that enables the user to obtain the number of compounds in the server’s database that are similar to the user’s target compound. Here, a fingerprint of compound is modeled as 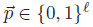 (*i.e.,* a bit string of length *ℓ*). An equivalent way to refer to 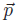 is the set of all indices *i* where *p_i_* = 1. We denote such a set by ***p***. The similarity of two compounds ***p***, ***q*** is then measured by *Tversky index* which is parameterized by *α, β >* 0 and is defined as:

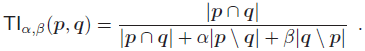

Tversky index is useful since it includes several important similarity measurements such as Jaccard Index (JI, which is exactly TI_1,1_ and also known as Tanimoto Index) and Dice index (which is exactly TI_1_*_/_*_2,1_*_/_*_2_) [22]. First, we introduce the basic idea and two efficient techniques for improving database privacy. Then, we describe our full proposed protocol.

### Basic idea

We firstly consider the simplest case that the user has (the fingerprint of) a target compound ***q*** as a query and the server’s database consists of only a single fingerprint ***p***. The case of a larger database is discussed later. The goal here is to detect whether or not the Tversky index of ***p*** and ***q*** is larger than a given threshold 1 ≥ *θ >* 0. The main idea of our approach is to calculate the score

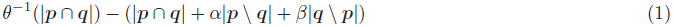

from *encrypted* fingerprints ***p*** and ***q*** by an additive-homomorphic cryptosystem. The score is non-negative if and only if the Tversky index of ***p*** and ***q*** is at least *θ*. Now since |***p*** \ ***q***| = |***p***| − |***p*** ∩ ***q***| and a similar relation holds for |***q*** \ ***p***|, the score (1) is positively proportional to

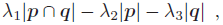

where *λ*_1_ = *c*(*θ*^−1^ − 1 + *α* + *β*), *λ*_2_ = *cα*, *λ*_3_ = *cβ* and any positive value *c*. We assume that the parameters and the threshold for the Tversky index are rational numbers denoted by *α* = *µ_a_/γ*, *β* = *µ_b_/γ* and *θ* = *θ_n_/θ_d_*, where *µ_a_*, *µ_b_*, *γ*, *θ_n_* and *θ_d_* are non-negative integers. By using *c* = *γ^θ^_n_g*^−1^ under this assumption, *λ*_1_, *λ*_2_ and *λ*_3_ become non-negative integers where *g* is the greatest common divisor of *γ*(*θ_d_* − *θ_n_*) + *θ_n_* (*µ_a_* + *µ_b_*), *θ_n_µ_a_* and *θ_n_µ_b_*.

Motivated by this observation, we define the following modified score, called the **threshold Tversky index**:

#### Definition 1.

*Given parameters α and β and a threshold θ for the Tversky index which are rational numbers denoted by α* = *µ_a_/γ, β* = *µ_b_/γ and θ* = *θ_n_/θ_d_ where µ_a_, µ_b_, γ, θ_n_ and θ_d_ are non-negative integers, then the* threshold Tversky index 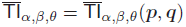 *for fingerprints* ***p*** *and* ***q*** *is defined by*

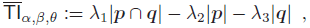

*and non-negative integer parameters λ*_1_*, λ*_2_ *and λ*_3_ *are defined by*

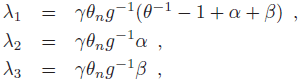

*where g is the greatest common divisor of γ*(*θ_d_* − *θ_n_*) + *θ_n_*(*µ_a_* + *µ_b_*)*, θ_n_µ_a_ and θ_n_µ_b_.*

By the above argument, we have TI*_α,β_*(***p***, ***q***) ≥ *θ* if and only if 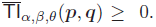. Therefore, the user can know whether or not his/her target compound ***q*** is similar (i.e., TI*_α,β_*(***p***,***q***) ≥ *θ*) to the fingerprint ***p*** in the database, by obtaining only the value 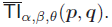.

In the protocol, the bits of the user’s target fingerprint ***q*** and the value |***p***| held by the server are both encrypted using the user’s public key. Since 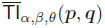 can be computed by the addition of these values and multiplication by integers, the protocol can calculate (without the secret key) a ciphertext of 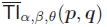, which is then decrypted by the user. For simplicity, we will abuse the notation and write TI(***p***,***q***), 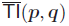 without subscripts *α, β, θ* when the context is clear.

We emphasize that our protocol does not use time-consuming cryptographic methods such as GP-MPC and FHE, and data transfer occurs only twice during an execution of the protocol. Hence, our protocol is efficient enough to scale to large databases.

### Database security enhancement techniques against regression attack

As discussed in Introduction section, the server needs to minimize returned information in order to minimize the success ratio of the regression attack. That is, the ideal situation for the server is that the user learns only the similarity/non-similarity property of fingerprints ***p*** and ***q***, without knowing any other information about the secret fingerprint ***p***. This means that only the sign of 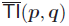 should be known by the user. However, in our basic protocol, the value of 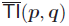 is fully obtained by the user; Database privacy is not protected from regression attacks. (See the Security analyses section for details.) In order to send only the sign of 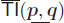, we firstly considered using a bit-wise decomposition protocol [23] for extracting and sending only the sign bit of 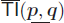. Although this approach is ideal in terms of security, the protocol requires more than 30 rounds of communications, which is much more efficient than using GP-MPC or FHE, but rather time-consuming for large-scale databases. Therefore, here we propose the novel technique of using dummy replies, which requires only one round of communication while sufficiently minimizing information leakage of ***p***. In the proposed technique, besides its original reply 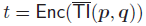, the server also chooses random integers *φ*_1_*,…,φ_n_* from a suitable interval and encrypts those values under the user’spublickey pk. Then the server sends the user a collection of ciphertexts *t,* Enc(*φ*_1_)*,…,* Enc(*φ_n_*) that are shuffled to conceal the true ciphertext *t*, as well as the number *s*_d_ of dummy values *φ_k_* with *φ_k_* ≥ 0. The user decrypts the received *n* + 1 ciphertexts, counts the number *s*_c_ of non-negative values among the decryption results, and compares *s*_c_ to *s*_d_. Now we have 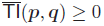 if and only if *s*_c_ − *s*_d_ = 1; therefore, the user can still learn the sign of 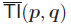, while the actual value of 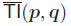 is concealed by the dummies. We have confirmed that the information leakage of ***p*** approaches zero as the number of dummies becomes large; see the Security analyses for pudding dummies section for more detailed discussion. (We have also developed another security enhancement technique using sign-preserving randomization of 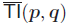; see Section 2 of Additional File 1 for details.)

### Database security enhancement technique against illegal query attack

Illegal query attacks can be prevented if the server can detect whether or not the user’s query is valid. To keep user privacy, the server must conduct this task without obtaining more information than the validity/invalidity of the query. In fact, this functionality can be implemented by using the NIZK proof by Sakai et al. [21] mentioned in the Non-interactive zero-knowledge proof section. The improved protocol requires the user to send the server a proof associated with the encrypted fingerprint bits *q_i_*, from which the server can check whether ***q*** is indeed a valid fingerprint (without obtaining any other information about ***q***); the server aborts the protocol if ***q*** is invalid. Here we use the “lifted” ElGamal cryptosystem as our basic encryption scheme to apply Sakai’s scheme. (We note that if we require the user to send `Enc`(−|***q***|) used by server’s computation, then another NIZK proof is necessary to guarantee the validity of the additional ciphertext, which decreases the communication efficiency of our protocol. Hence our protocol requires the server to calculate `Enc`(−|***q***|) by itself.) The formal definition of the valid query is given in the Database privacy in malicious model section.

### Secure similar compounds counter

For the general case that the database consists of more than one fingerprint ***p***, we propose the protocol shown in Algorithm 1 to count the number of fingerprints ***p*** similar to the target fingerprint ***q***. In the protocol, the server simply calculates the encryption of the threshold Tversky indices for all database entries and, as discussed above, replies with a shuffled collection of these true ciphertexts and dummy ciphertexts, as well as the number *s*_d_ of nonnegative dummy values. Then the value *s*_c_−*s*_d_ finally obtained by the user is equal to the number of similar fingerprints ***p*** in the database.

### Parameter settings of the protocol

Decrypting an encrytion of too large value needs huge computation cost if the lifted-ElGamal cryptosystem is used. Therefore, in order to keep the consistency and efficiency of the protocol, the range of 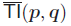 should not be too large. i.e., the integer parameters *λ*_1_, *λ*_2_ and *λ*_3_ in the threshold Tversky index should not be too large. In fact, this will not cause a problem in practice; For example, the parameters become *λ*_1_ = 9, *λ*_2_ = *λ*_3_ = 4 for computing 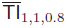 which is a typical setting of a chemical compound search. In this case, a minimum value and a maximum value of 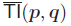 is -664 and 166 for 166 MACCS keys, which is a sufficiently small range. (See Section 3 of Additional File 1 for details.)

#### Algorithm 1 The secure similar compounds counter (SSCC)

- Public input: Length of fingerprints *ℓ* and parameters for the Tversky index *θ* = *θ_n_/θ_d_, α* = *µ_a_/γ, β* = *µ_b_/γ*
- Private input of a user: Target fingerprint ***q***
- Private input of a server: Set of fingerprints *P* = {***p***^(1)^*,…,* ***p***^(^*^M^*^)^}

1. (Key setup of cryptosystem) The user generates a key pair (pk, sk) by the key generation algorithm KeyGen for the additive-homomorphic cryptosystem and sends public key pk to the server (the user and the server share public key pk and only the user knows secret key sk).
2. (Initialization) The user encrypts his/her fingerprint ***q*** as a vector of ciphertexts: 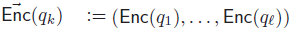. He/she also generates ***v*** as a vector of proofs. Each proof *v_i_* is associated with `Enc`(*q_i_*).
3. (Query of entry) The user sends the vector of ciphertexts 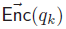 and the vector of proofs ***v*** to the server as a query.
4. (Query validity verification) The server verifies the validity of 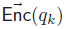 by testing the vector of proof ***v***. If ***v*** does not pass the server’s test, the user cannot move on to the next step.
5. (Calculation of threshold Tversky index)

a. The server calculates the greatest common divisor of *γ*(*θ_d_* − *θ_n_*) + *θ_n_*(*µ_a_* + *µ_b_*), *θ_n_µ_a_* and *θ_n_µ_b_* as *g*, and calculates *λ*_1_ = *γθ_n_g*^−1^ (*θ*^−1^ − 1 + *α* + *β*), *λ*_2_ = *γθ_n_g*^−1^*α,* and *λ*_3_ = *γθ_n_g*^−1^*β.*
b. The server calculates 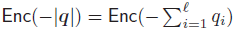 from 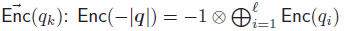.
c. **for** *j* = 1 to *M* **do**

i. The server calculates 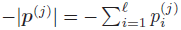 and encrypts it to obtain a ciphertext `Enc`(−|***p***^(^*^j^*^)^|).
ii. The server calculates a ciphertext *t_j_* of threshold Tversky index 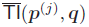.

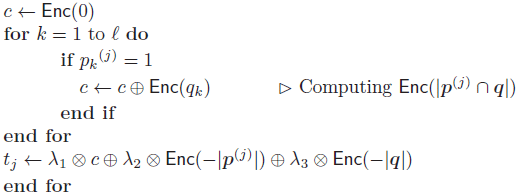
6. (Padding of dummies)

a. The server generates a set of dummy values {*φ*_1_*,…,φ_n_*} and counts the number *s*_d_ of non-negative dummies *φ_i_* ≥ 0.
b. The server encrypts *φ_i_* to obtain a ciphertext `Enc`(*φ_i_*) for *i* = 1*,…,n*.
c. The server shuffles the contents of the set *T* = {*t*_1_*,…,t_M_,* `Enc`(*φ*_1_)*,…,* `Enc`(*φ_n_*)}.
7. (Return of matching results) The server sends *T* and *s*_d_ to the user.
8. (Decryption and counting) The user decrypts the contents of *T* and counts the number *s*_c_ of non-negative values.
9. (Evaluation) The user obtains *s*_c_ − *s*_d_ as the number of similar fingerprints in the database.

## Security analyses

In this section, we evaluate security of SSCC by several approaches.

In the area of cryptology, the following two standard security models for two-party computation have been considered:

- *Semi-honest model*: Both parties follow the protocol, but an adversarial one attempts to infer additional information about the other party’s secret input from the legally obtained information.
- *Malicious model*: An adversarial party cheats even in the protocol (e.g., by inputting maliciously chosen invalid values) in order to illegally obtaining additional information about the secret.

We analyze user privacy and database privacy in both the semi-honest and malicious models. For the database privacy, we firstly compare attack success ratios for the case of using our method which aims to output a binary sign and the other case of using the previous methods which aim to output a similarity score, and show that outputting a binary sign improves database privacy. We also evaluate security strength of our method against a regression attack by comparing attack success ratios for the case of using dummies and the ideal case that uses a versatile technique (such as GP-MPC and FHE) to output a binary sign, and show that the security strength for the case of using dummies is almost the same as the ideal case under realistic settings.

### User privacy

The semantic security of the encryption scheme used in the protocol (see the Additively homomorphic encryption scheme section) implies immediately that the server cannot infer any information about the user’s target fingerprint ***q*** during the protocol. This holds in both the semi-honest and malicious models.

### Thresholding largely improves database privacy

We mentioned in the introduction section that minimizing information returned from the server reduces success ratio of regression attack. Therefore, SSCC aims for “ideal” case in which the user learns only the sign of 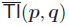 during the protocol. The previous methods that compute Jaccard Index aim for the “plain” case, in which the user fully learns the value TI(***p***,***q***). Here we evaluate the efficiency of the thresholding by comparing success probabilities of regression attack for those two cases. We consider the general case in which the user is allowed to send more than one query and those queries are searched by Jaccard Index. We also suppose that the database consists of a single fingerprint ***p*** in order to clarify the effect of thresholding.

The goal of an attacker is to reveal ***p*** by analysing the results returned from the server. It is generally effective for the attacker to exploit the difference between the two outputs obtained by sending two different queries. In fact, when the server returns 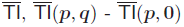 becomes positive if and only if *p_i_* = 1, where ***q*** = (0*,…,q_i_* = 1*,…,* 0) and **0** = (0*,…,* 0). This means that the attacker can reveal any bit in ***p*** by sending the single query after sending the first query **0**. Therefore, ***p*** can be fully revealed by sending only *ℓ* + 1 queries. On the other hand, there is no deterministic attack for revealing ***p*** from only the sign of 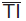, because two different inputs do not always lead to different outputs. Since we know of a linear algorithm that fully reveals ***p*** in response to at most 2*ℓ* queries after making a “hit” query ***q*** such that 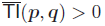, here we evaluate database privacy by the probability of making at least one hit query when the user is allowed to send *x* queries. (See Section 4 of Additional File 1 for details.) This probability is denoted as

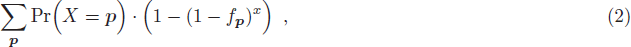

where *f****_p_***, defined as follows, is the probability that the user makes one hit query with a single trial when ***p*** is given.

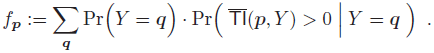

For ease of calculation, we computed the upper bound of equation (2) for *x* = 1,10,10^2^,*…,* 10^6^ and *θ* = 0.7,0.8,0.9,1.0. (See Section 5 of Additional File 1 for details.) Since publicly available 166 MACCS keys are the most popular fingerprint for chemical compound searches, we set *ℓ* to 166. From the results shown in Fig. 3, we can see that the probability of making a hit query is sufficiently small for practical use even though the user is allowed to send a million of queries. Considering that the user learns ***p*** by using no more than *ℓ* + 1 queries when he/she learns 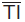, we can conclude that database privacy is dramatically improved by thresholding. In other words, the proposed protocol, which aims to output only the sign of the similarity score, has stronger security than other previous methods, which directly output similarity scores.

**Figure 3:**
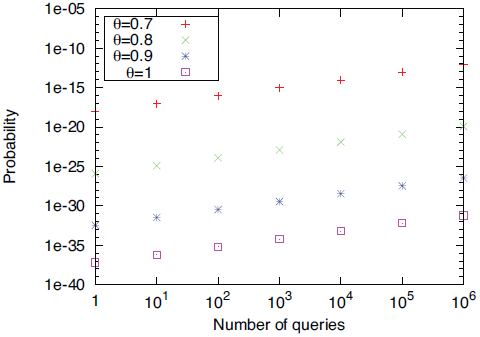
Upper bounds of the probabilities that the user has at least one hit query out of making 1, 10*,…,* 10_6_ queries. Note that the hit query becomes the critical hint for revealing database information. Each line shows the results with one of the four different thresholds.

### Security analyses for padding dummies

We showed that the output privacy in the “ideal” case is significantly improved from the “plain” case. Here we experimentally evaluate how the actual situation of our proposed protocol is close to the “ideal” case.

Before going into detail analyses, let us discuss how to generate dummies. It is ideal for the server privacy to generate a dummy according to the same distribution where 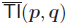 is generated from. However, this is not realistic because 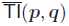 is determined by both ***p*** and ***q*** which is user’s private information. Therefore, in our analyses, we assume that a dummy is generated from uniform distribution over possible values of 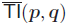. For example, if possible values of 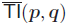 is {1,2,3,4,5}, dummies are randomly selected from any one of them. The purpose of padding dummies is to mitigate the risk of leaking 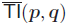. In order to clarify the effect of the use of dummy values, we concentrate on the basic case; the database contains a single ***p***, and there exist *k* possible values of 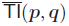. *i*-th value of the *k* possible values arises as the true 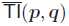 according to the probability *w_i_*. Namely, true 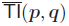 is generated from the multinomial distribution with *k* different probabilities ***w*** = *w*_1_*,…,w_k_*, while dummies are generated from the multinomial distribution with equal probability 1*/k*. To conduct stringent analyses, we assume that the user knows ***w***, and he/she also knows that dummies are uniformly distributed over *k* possible 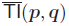.

The security provided by our protocol can be formalized in the following manner. First we recall that, in our protocol, the server computes encryption of 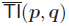 and encryption of dummy values *φ*_1_,…,*φ_n_*, and then sends the user the n + 1 encrypted values as well as the number of positive dummy values in *φ*_1_,…,*φ_n_*. For the purpose of formalizing the security, we introduce a “fictional” server that performs the following: It first receives the encrypted values 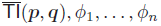 from the real server. Secondly, it gets the sign of 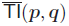. (We note that a real server cannot do it since it requires unrealistic computational power that breaks the security of the encryption scheme, so this is just fictional for the sake of mathematical definition.) Thirdly, it generates another dummy value 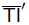 randomly, and independently of the values of 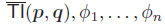 (except for the sign of 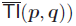, in the following manner:

- If 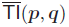 is positive, then 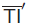 is chosen randomly from positive values.
- If 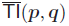 is negative, then 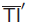 is chosen randomly from negative values.

Finally, the fictional server sends the user an encryption of 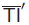 (instead of 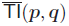) as well as the encrypted *φ*_1_*,…,φ_n_* and the number of positive values in *φ*_1_*,…,φ_n_*. We note that, when the user receives a reply from the fictional server, the user can know the sign of 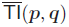 which is the same as that of 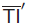, but cannot know any other information on 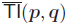 since 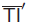 is independent of 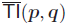. In the setting, the following property can be proven:

#### Theorem 1.

*Suppose that the user cannot distinguish, within computational time* TIME*, the sets of decrypted values of ciphertexts involved in outputs of the real server and of the fictional server. Then any information computable within computational time* TIME *from the decryption results for output of the real server is equivalent to information computable within computational time* TIME′ *from the sign of* 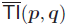 *only, where* TIME′ *is a value which is close to* TIME.

*Proof.* Let 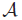 be an algorithm, with running time TIME, which outputs some information on the decrypted values for an output of the real server. We construct an algorithm 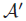 which computes, from the sign of 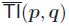 only, an information equivalent to the information computed by 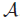. The construction is as follows; from the sign of 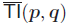, 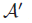 generates dummy values by mimicking the behavior of the fictional server, and then 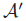 inputs these dummy values to a copy of 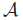, say 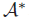, and gets the output of 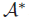. Now if the output of 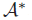 is not equivalent to the output of 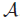, then the definition of 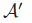 implies that the probability distributions of the outputs of 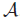 with inputs given by the decrypted values for outputs of the real server and of the fictional server are significantly different (since 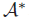 used in 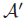 is a copy of 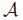); it enables the user to distinguish the two possibilities of his/her received values by observing the output of 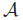, but this contradicts the assumption of the theorem. Therefore, the output of 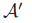 is equivalent to the output of 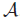 as claimed. Moreover, the computational overhead of 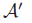 compared to 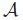 is just the process of generating dummy values by mimicking the behavior of the fictional server; it is not large (i.e., TIME′ is close to TIME as claimed) since the server-side computation of our proposed protocol is efficient. Hence, the theorem holds.

Roughly rephrasing, if the assumption of the theorem is true for a larger TIME, then the actual situation of our proposed protocol becomes closer to the “ideal” case provided we focus on any information available from efficient computation. As a first step to evaluate how the assumption is plausible (i.e., how the value TIME in the assumption can be large), we performed computer experiments to show that some natural attempts to distinguish the actual and the fictional cases do not succeed, as explained below.

In this experiment, we evaluate the security of our protocol by comparing the probabilities that the user correctly guesses the value 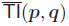 in two cases: The case in which the user makes a guess based only on a prior knowledge ***w***, and the other case in which the user makes a guess based on the observation of the search result under the condition that he/she knows ***w***.

For the first case, the user’s best strategy for guessing 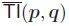 is to choose the *i*_0_-th possible value, where

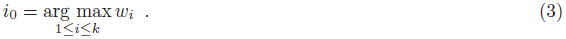

In this case, the success probability of the guess is *w_i_*_0._

Let us consider the best strategy for the second case. As described above, we consider an practical case that *n* dummy values *φ*_1_*,…,φ_n_* chosen from the *k* possible values uniformly at random, and the user makes a guess from the received *n* + 1 shuffled values *φ*_1_*,…,φ_n_,* 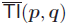. Now suppose that the user received the *i*-th possible value *a_i_* times for each 1 ≤ *i* ≤ *k* (hence 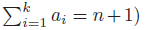). Since the choices of *φ*_1_*,…,φ_n_* are independent of 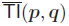, the probability that the user received *i*-th possible value *a_i_* times for each 1 ≤ *i* ≤ *k* and that 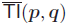 is *i*_0_-th possible value is

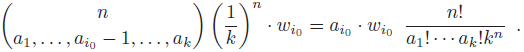

Therefore, the conditional probability that 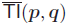 is the *i*_0_-th possible value, conditioned on the set of the user’s received values, is

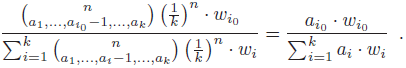

This implies that the user’s best strategy is to guess that 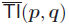 is the *i*_0_-th possible value, where

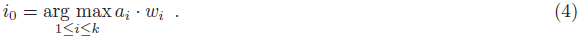

We estimated success probabilities of user’s guess for the both cases by simulation experiments. Here we assumed typical case when TI_1,1,0_*_._*_8_ and 166 MACCS keys are used. In this case, *k* = 831 and we performed the experiments for *n* = 831 × 10^0^, 831 × 10^1^,*…,* 831 × 10^4^ on three different distributions of 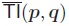 which were obtained by the following schemes:

1. We randomly selected one fingerprint ***q*** from ChEMBL and calculated 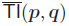 for all the entries in ChEMBL and used the observed distribution as ***w***. In our experiment, 177159-th fingerprint was selected as ***q*** (referred as ***W***^ChEMBL–177159^).
2. The same scheme as 1) was used when ***q*** was 265935-th fingerprint (referred as ***w***^ChEMBL–265935^).
3. We randomly selected a value from 1*,…,k* for *m* times and count frequency of *i* as *h_i_* and set *w_i_* = *h_i_/m* (referred as *w*^random^). We used *k* × 5 as *m*.

All the distributions used here are shown in Section 6 of Additional File 1.

We performed 100,000 trials for each experiment. Each trial consisted of choosing *φ*_1_*,…,φ_n_* uniformly at random; choosing 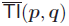 according to ***w***; deciding the user’s guess *i*_0_ by formula (3) and formula (4) respectively (we adopted a uniformly random choice if there were more than one such *i*_0_); and checking whether or not 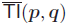 was the *i*_0_-th possible value for both rules (i.e., the user’s guess succeeded). The results of the experiment are given in Table 1; they show that the user’s attack success probability became significantly close to the ideal case when a sufficiently large number of dummies were used; therefore, our technique of using dummies indeed improves the output privacy.

**Table 1:** The experimental success ratios of the user’s guess based on the server’s return and the prior distribution of true value (*n* = 813*,…,* 813 × 10^4^), and success probability based only on a guess from the prior distribution (ideal value). 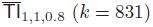 is assumed and results are calculated for five different numbers of dummies (*n* = 831,831× 10^1^, 831 ×10^2^, 831 × 10^3^, 831 × 10^4^) are used for three different distributions: ***w***^ChEMBL–177159^ w^ChEMBL–265935^ are actual distributions of 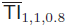 on ChEMBL obtained by querying two randomly selected fingerprints from ChEMBL, ***w***^rand^ is obtained by randomly selecting a value from 1*,…,k* for *m* = 5 × 831 times and dividing each observed frequency by *m*.

| $n$ | 831 | $831 \times 10^1$ | $831 \times 10^2$ | $831 \times 10^3$ | $831 \times 10^4$ | Ideal value |
| --- | --- | --- | --- | --- | --- | --- |
| $\mathbf{w}^{\text{ChEMBL-177159}}$ | 0.03552 | 0.01738 | 0.01101 | 0.01009 | 0.00977 | 0.00981 |
| $\mathbf{w}^{\text{ChEMBL-265935}}$ | 0.02991 | 0.01337 | 0.00903 | 0.00798 | 0.00784 | 0.00807 |
| $\mathbf{w}^{\text{rand}}$ | 0.00914 | 0.0041 | 0.00309 | 0.00279 | 0.00305 | 0.00289 |

### Security analyses for padding dummies for the case when the user is allowed to send more than one query

One might suspect that the attacker can detect the true 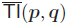 by sending the same query twice and finding the value which is appeared in both results. However, this attack does not easily succeed if *n* is sufficiently larger than *k* (i.e., ideally, all possible values of 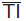 are covered by sufficient number of dummies), and we consider that *k* is not too large in practice as we discussed in Parameters settings of the protocol section.

In order to evaluate the security achieved by the padding dummies for the case when the user is allowed to submit *L* queries, we performed following analyses. Here we evaluate the security achievement by comparing the case of using our protocol based on the padding dummy and the ideal case of returning only the sign of 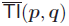. In order to perform rigorous analyses, we assume the most severe case in which the attacker keeps sending the same query *L* times. For this case, the probability that 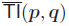 is the *i*_0_-th possible value after sending *L* queries on condition that frequency of *i*-th possible value of *j*-th query 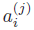 for *j* = 1*,…, L* is

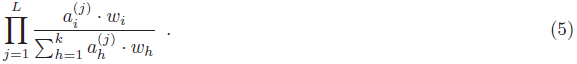

This implies that the user’s best strategy is to choose *i*-th possible value which maximizes equation (5). As mentioned above, we compared the success ratio of the attack based on the above strategy and the ideal success ratio when the user makes the guess only from the given distribution ***w***. We also assumed more realistic case that user did not know the exact distribution of dummy but knew the distribution that was similar to the actual distribution the server used. For the evaluation of this case, we generated dummies from the distribution ***u***, which was slightly different from uniform distribution, while the user assumed that dummies were generated from uniform distribution. ***u*** was generated as follows:

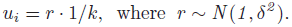

We performed the experiment for *L* = 1,10,10^2^,*…,* 10^5^, *n* = 831 × 10,831 × 50,831 × 10^2^ and *δ* = 0,0.05, 0.1,0.15, 0.2 based on the same approach used in the evaluation of single query security. i.e., for each trial, *n* dummies were randomly chosen according to ***u*** (note that ***u*** was equal to uniform distribution when *δ* = 0), true value 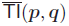 was selected according to ***w*** and the attacker’s guess was made based on the equation (5). We performed 10,000 trials for each triplet of *L*, *n* and *δ*. Those experiments were conducted for the same three distributions: ***w***^ChEMBL–177159^, ***w***^ChEMBL–265935^ and ***w***^ChEMBL–random^. We compared the success ratio of the attack and the ideal success ratio when the user made the guess without seeing search results. The results are shown in Fig. 4. The success ratio of user’s attack decreased as the number of dummies increased and became closer to the ideal value when the sufficient number of dummies are given, even for the case that a large number of queries were sent. Although an efficient method for dummy generation remains as a future task, the results also show that hiding the distribution of dummy is significantly effective for protecting database privacy and the user has to know it with high accuracy in order to steal extra information from the server.

**Figure 4:**
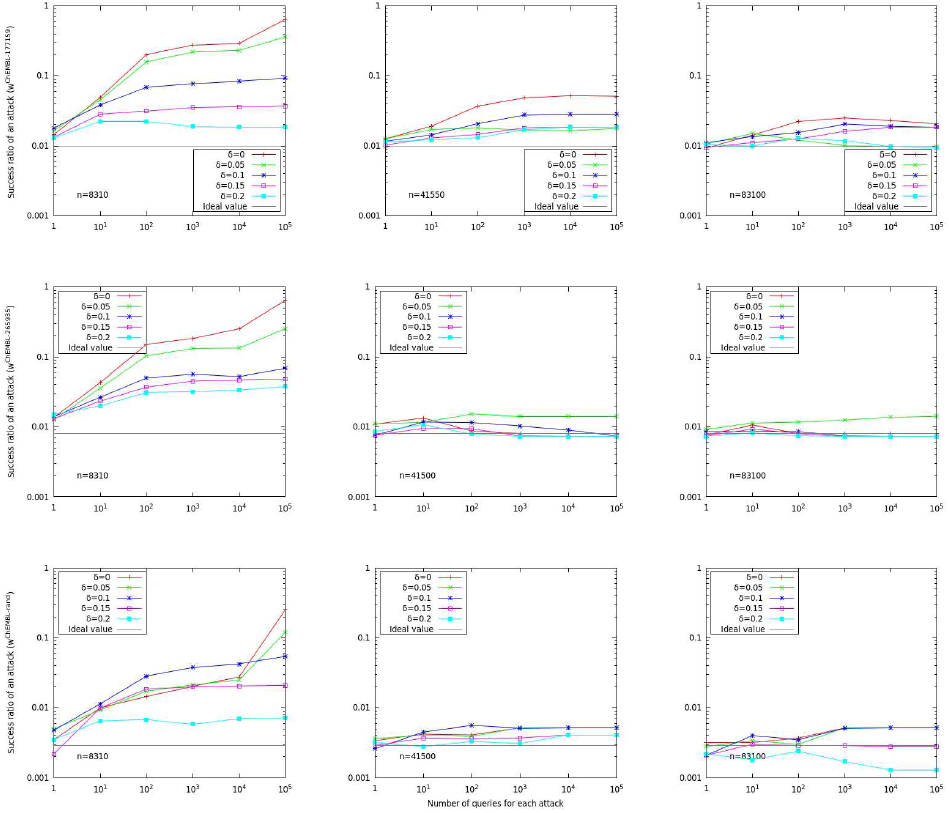
The comparison of the experimental success ratios of the user’s guess based on the server’s return as well as the prior distribution of true value when the user sends many queries (*δ* = 0*,…,* 0.2), and success probability based only on a guess from the prior distribution (ideal value). 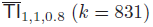 is assumed and results are calculated for three different numbers of dummies (*n* = 831 × 10, 831 × 50, 831 × 10^2^) when the user sends *L* = 1,10*,…,* 10^5^ queries and three different distributions: ***w***^ChEMBL–177159^ and ***w***^ChEMBL–265935^ are actual distributions of 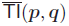 on ChEMBL obtained by querying two randomly selected fingerprints from ChEMBL, ***w***^rand^ is obtained by randomly selecting a value from 1*,…,k* for *m* = 5 × 831 times and dividing each observed frequency by *m*.

### Database privacy in malicious model

For our protocol, the difference between the malicious and semi-honest models is that in the malicious model the user may use an invalid input ***q*** whose components *q_i_* are not necessarily in {0, 1}. If the user chooses ***q*** in such a way that some component *q_i_* is extremely large and the remaining *ℓ* − 1 components are all zero, then 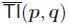 will also be an extreme value (distinguishable from the dummy values) and depend dominantly on the bit *p_i_*; therefore, the user can almost surely guess the secret bit *p_i_*. Since our protocol detects whether or not *q_i_* is a bit value without invading user privacy, it can safely reject illegal queries and prevent any illegal query attacks, including above case.

## Performance evaluation

In this section, we evaluate the performance of the proposed method on two datasets created from ChEMBL.

### Implementation

We implemented the proposed protocol using the C++ library of elliptic curve ElGamal encryption [24], in which the NIZK proposed in the previous study [21] is also implemented.

For the implementation, we used parameters called secp192k1, as recommended by SECG (The Standards for Efficient Cryptography Group). These parameters are considered to be more secure than 1024-bit RSA encryption, which is the most commonly used public-key cryptosystem. The implementation of

Owing to the limitation of the range of plaintext, the implementation here does not include sign-preserving randomization. For the purpose of comparison, we also implemented a GP-MPC protocol by using Fairplay [25]. In order to reduce the circuit size of the GP-MPC, we implemented s simple task that computes the sign of Tversky index between a query and a fingerprint in the database, and repeated the task for all the fingerprints in the database. Thus the CPU time and data transfer size of the implementation is linear to the size of database.

### Experimental setup

The Jaccard index along with the threshold *θ* = 0.8 were used for both protocols. For SSCC, we used 10,000 dummies. These two implementations were tested on two datasets: one, referred to as ChEMBL_1000, was the first 1000 fingerprints stored in ChEMBL, and the other, referred to as ChEMBL_Full, was 1,292,344 fingerprints in the latest version of ChEMBL. All the programs were run on a single core of an Intel Xeon 2.9 GHz on the same machine equipped with 64 GB memory. To avoid environmental effects, we repeated the same experiment five times and calculated average values.

### Results

The results are shown in Table 2. Despite the proposed method including elaborate calculation like the NIZK proof, we can see from the results that both the CPU time and communication size of the proposed method are significantly smaller than those of the GP-MPC protocol. Furthermore, it is clear that SSCC provides industrial-strength performance, considering that it works, even on a huge database like ChEMBL_Full, taking no more than 167 s and 173 s for the server and client respectively.

**Table 2:** CPU time and communication size of secure similar compounds counter (SSCC) and those of general-purpose multi-party computation (GP-MPC). The experiment on ChEMBL_Full by GP-MPC did not finish within 24 hours.

|  | ChEMBL_1000 | ChEMBL_Full |
| --- | --- | --- |
| CPU time (s) |  |  |
| SSCC (server) | 0.69 | 167.19 |
| SSCC (client) | 1.53 | 172.37 |
| GP-MPC (server) | 4,075.15 | — |
| GP-MPC (client) | 4,366.18 | — |
| Communication size (MB) |  |  |
| SSCC (server → client) | 2.24 | 265.33 |
| SSCC (client → server) | 0.03 | 0.03 |
| GP-MPC (server → client) | 42.50 | — |
| GP-MPC (client → server) | 2,128.00 | — |

The experiment on ChEMBL_Full by using GP-MPC did not finish within 24 hours. Since both CPU time and communication size are exactly linear to the size of database for the GP-MPC protocol, the results of ChEMBL_Full for GP-MPC are estimated to be more than 1600 hours for both sides and 3 Gbyte data transfer from client to server, considering the results of ChEMBL_1000.

By using simple data parallelization, the computational speed will be improved linearly with the number of CPUs. Since all the programs were run on the same machine there was almost no latency for the communication between the two parties in these experiments. Therefore, GP-MPC, whose communication size is huge, is expected to require far more time when it runs on an actual network that is not always in a good condition. The other important point is that SSCC requires only two data transfers, which enables data transfer after off-line calculation. On the other hand, GP-MPC must keep online during the search because of the high communication frequency. We also note that it took less than 100 MB to compile SSCC, while GP-MPC required more than 16 GB. Considering these observations, SSCC is efficient for practical use. It is known that several techniques improve the performance of GP-MPC and the previous work by Pinkas et al. [26] reported that Free XOR [27] and Garbled Row Reduction [26], which are commonly used in state-of-the-art GP-MPC methods [28] [29] [30] [31], reduced running time and communication size by factors of 1.8 and 6.3 respectively when a circuit computing an encryption of AES was evaluated. Though these techniques are not implemented in Fairplay, we consider that GP-MPC is yet far less practical for the large-scale chemical compound search problem compared to our method which improved running time and communication size by factors of 36, 900 and 12, 000.

## Conclusion

In this study, we proposed a novel privacy-preserving protocol for searching chemical compound databases. To our knowledge, this is the first practical study for privacy-preserving (for both user and database sides) similarity searching in the fields of bioinformatics and chemoinformatics. Moreover, the proposed method could be applied to a wide range of life science problems such as searching for similar single-nucleotide polymorphism (SNP) patterns in a personal genome database. While the protocol proposed here focuses on searching for a number of similar compounds, we are examining further improvements of the protocol such as the client being able to download similar compounds; we expect this on-going study to further contribute to the drug screening process. In recent years, open innovation has been attracting attention as a promising approach for speeding up the process of new drug discovery [32]. For example, research on neglected tropical diseases including malaria has been promoted by the recent attempt to share chemical compound libraries in the research community. In spite of high expectations, such an approach is still limited to economically less important problems on account of privacy problems [33]. Therefore, privacy-preserving data mining technology is expected to be the breakthrough promoting open innovation and we believe that our study will play an important role.

## Acknowledgement

KS thanks Yusuke Sakai and Takahiro Matsuda for fruitful discussions.

## A An Encryption Scheme with Zero-Knowledge Proof

For self-containment, we briefly recall the lifted ElGamal encryption scheme with the non-interactive zero-knowledge proof (NIZK) system which proves that the plaintext is either 0 or 1, proposed in [21], and provide some intuition for it. Let 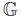 be a group of prime order *p* such that the Decisional Diffie-Hellman assumption holds (i.e., the resulting encryption scheme provides a standard security property called semantic security). The public key is (*g, h, f*) where *g, f* are random generators in 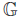 and *h* = *g^z^* for *z* randomly picked from 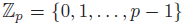. The secret key is *z*. Let *H* be a hash function. Encrypting a plaintext *b* ∈ {0,1} gives a ciphertext (*C*_1_ = *g^u^,C*_2_ = *h^u^f^b^*), for random 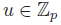. Decryption computes 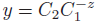 and outputs 1 if *y* = *f* or outputs 0 if *y* = 1.

- Encrypting a plaintext 0 outputs the ciphertext (*C*_1_*, C*_2_)where

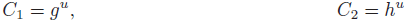

where *u* is randomly picked from 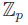. The proof that its plaintext is either 0 or 1 consists of

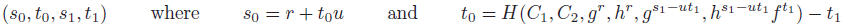

where *r, s*_1_*, t*_1_ are randomly picked from 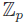.
- Encrypting a plaintext 1 outputs the ciphertext (*C*_1_*, C*_2_) where

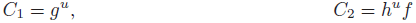

where *u* are randomly picked from 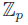. The proof that its plaintext is either 0 or 1 consists of

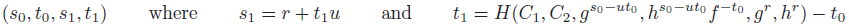

where *r, s*_0_*, t*_0_ are randomly picked from 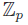.

The verification of the proof is done by computing

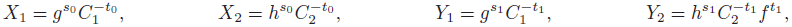

and checking whether

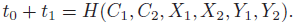

Correctness can be verified straightforwardly. As for security, we provide the intuition as follows:

- **Soundness.** The relations for *X*_1_*, X*_2_ prove that the plaintext is 0, while the relations for *Y*_1_*, Y*_2_ prove that the plaintext is 1. The prover can produce exactly one real proof and one simulated fake proof due to the fact that the term *t*_0_ + *t*_1_ is constrained to the hash of *r*. In case of encrypting a bit *b*, the prover chooses 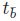 freely, where 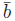 is the flipped bit of *b*. The fact that the prover cannot control *t_b_* implies that it has to know the ciphertext randomness *u* in order to produce the correct *s_b_* (*à la* the Fiat-Shamir Paradigm [34]). This implies that there exists *u* so that (*C*_1_*, C*_2_) forms an encryption of *b*, and hence the soundness. In other words, since the output of the hash function *H* cannot be predicted before determining its input, in order to satisfy the condition *t*_0_ + *t*_1_ = *H*(*C*_1_*, C*_2_*, X*_1_*, X*_2_*, Y*_1_*, Y*_2_), the only possible strategy is to choose at least one of *t*_0_ and *t*_1_ in a consistent manner *after the input for H is determined*. Now note that

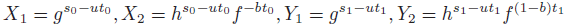

if (*C*_0_*, C*_1_) is a ciphertext of *b*. The shapes of *X*_1_ and *X*_2_, in particular the exponent −*bt*_0_ of *f* in *X*_2_, imply that *t*_0_ is uniquely determined from *X*_1_ and *X*_2_ *unless b* = 0. Similarly, the shapes of *Y*_1_ and *Y*_2_, in particular the exponent (1 − *b*)*t*_1_ of *f* in *Y*_2_, imply that *t*_1_ is uniquely determined from *Y*_1_ and *Y*_2_ *unless b* = 1. Therefore, *unless b* = 0 *or b* = 1, there is no room to adjust the value of *t*_0_ or *t*_1_ after the input for *H* is determined as required above. By the contraposition, the existence of a valid proof implies that the plaintext *b* must be 0 or 1, ensuring the soundness.
- **Zero-Knowledge.** Observe that, in both cases, the proof elements (*s*_0_*, t*_0_*, s*_1_*, t*_1_) distribute identically, due to the uniform randomness of *u, r,* 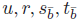 (in case of encrypting *b*). Hence, the information on the plaintext bit is hidden from the proof.

## B Further security enhancement technique by using sign-preserving randomization

As mentioned at the end of the Database security enhancement techniques against regression attack section, we can further improve the output privacy by using “sign-preserving randomization” of the threshold Tversky index. Namely, the server calculates 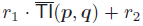 with random integers 0 ≤ *r*_2_ *< r*_1_ for each ***p*** and uses it instead of the true value 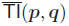 (the range of the dummy values *φ*_1_*,…,φ_n_* should also be changed accordingly). This modification makes it more difficult for the user to guess the true 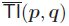; it does not affect the correctness of the protocol, since 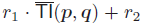 has the same sign as 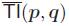. Therefore, it is qualitatively obvious that the output privacy is improved by the sign-preserving randomization. However, it is hard to quantitatively evaluate the achieved improvement; more detailed evaluations will be a future research topic.

## C Ranges of 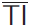 for typical parameter settings

In the Parameter settings of the protocol section, we mentioned that the range of 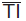 is not too large for practical cases of chemical compound search. Here we list the ranges of typical parameter settings in Table 3. In our protocol, it is sufficient to check whether or not the 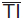 is positive value. Therefore, the range which is necessary to be verified is smaller in practical.

**Table 3:** Range of 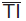 for the most typical parameter settings.

|  |  |  |  |  |  |
| --- | --- | --- | --- | --- | --- |
| 166 | 1.0 | 1.0 | 0.7 | ( 498 , -1162 ) | 499 |
| 166 | 1.0 | 1.0 | 0.8 | ( 166 , -664 ) | 167 |
| 166 | 1.0 | 1.0 | 0.9 | ( 166 , -1494 ) | 167 |
| 166 | 1.0 | 1.0 | 1.0 | ( 0 , -166 ) | 1 |
| 166 | 0.5 | 0.5 | 0.7 | ( 996 , -1162 ) | 997 |
| 166 | 0.5 | 0.5 | 0.8 | ( 166 , -332 ) | 167 |
| 166 | 0.5 | 0.5 | 0.9 | ( 332 , -1494 ) | 333 |
| 166 | 0.5 | 0.5 | 1.0 | ( 0 , -166 ) | 1 |
| 166 | 1.0 | 0.0 | 0.7 | ( 498 , -1162 ) | 499 |
| 166 | 1.0 | 0.0 | 0.8 | ( 166 , -664 ) | 167 |
| 166 | 1.0 | 0.0 | 0.9 | ( 166 , -1494 ) | 167 |
| 166 | 1.0 | 0.0 | 1.0 | ( 0 , -166 ) | 1 |
| 960 | 1.0 | 1.0 | 0.7 | ( 2880 , -6720 ) | 2881 |
| 960 | 1.0 | 1.0 | 0.8 | ( 960 , -3840 ) | 961 |
| 960 | 1.0 | 1.0 | 0.9 | ( 960 , -8640 ) | 961 |
| 960 | 1.0 | 1.0 | 1.0 | ( 0 , -960 ) | 1 |
| 960 | 0.5 | 0.5 | 0.7 | ( 5760 , -6720 ) | 5761 |
| 960 | 0.5 | 0.5 | 0.8 | ( 960 , -1920 ) | 961 |
| 960 | 0.5 | 0.5 | 0.9 | ( 1920 , -8640 ) | 1921 |
| 960 | 0.5 | 0.5 | 1.0 | ( 0 , -960 ) | 1 |
| 960 | 1.0 | 0.0 | 0.7 | ( 2880 , -6720 ) | 2881 |
| 960 | 1.0 | 0.0 | 0.8 | ( 960 , -3840 ) | 961 |
| 960 | 1.0 | 0.0 | 0.9 | ( 960 , -8640 ) | 961 |
| 960 | 1.0 | 0.0 | 1.0 | ( 0 , -960 ) | 1 |

## D The attack algorithm by using a hit query

In this section, we showed the full algorithm to reveal the server’s fingerprint when a “hit” query is given, as mentioned in the Thresholding largely improves output privacy section.

Here we assume following case: The user is allowed to send more than one query and the database consists of a single fingerprint ***p***. For each trial, the user learns from the server whether or not 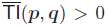. Prior to his/her attack, the user is given a hit query 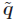 such that 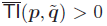. The goal of the algorithm shown here is to reveal the server’s fingerprint ***p***. Here, we denote the reverse bit of *x* as Rev(*x*).

### Theorem 2.

*Let there be bit vectors* ***p*** *and* 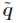 *where* 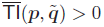 *holds for* ***p*** *and* 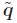*, and a bit vector* ***q*** *which is defined by* 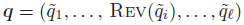*. Then the inequality* 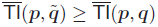 *holds if and only if* 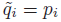.

*Proof.* If 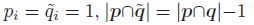 and 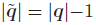. Therefore 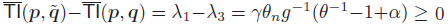 from the definition of 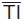 where 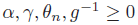 and 1 ≥ *θ>* 0.

If 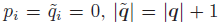. Therefore 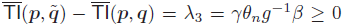 from the definition of 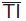 where *γ,θ_n_,****g*** ^−1^*,β* ≥ 0.

According to Theorem 2 and considering that *θ* > 0.5 is used for similarity search, it is trivial that 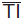 mostly keeps decreasing along with bits of 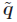 being reversed sequentially and finally reaches a minus value. Therefore, we can find a bit vector 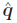 such that it satisfies

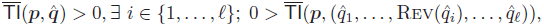

and we call the bit vector “detective” query.

The inequality 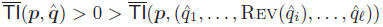 holds if and only if 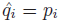 according to Theorem 2. Therefore the attacker knows whether or not 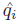 is equal to the database’s fingerprint *p_i_* from the observation of 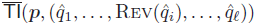. We show full description of the attack algorithm in Algorithm 2.

## E Derivation and calculation of upper bound of the probability for making at least one hit query

Here we provide a detailed derivation of the upper bound of (2) in the Thresholding largely improves database privacy section. In order to simplify the problem, the condition 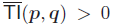 is replaced by the equal condition 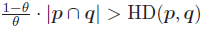 where HD(***p***,***q***) is the Hamming distance of ***p*** and ***q***. For further simplification, we use the bound 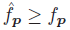, derived from |***p***|≥|***p*** ∩ ***q***|, where

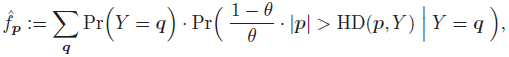

to obtain the following probability, which is the upper bound of the probability (2).

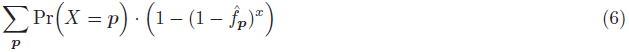

We use the known approximation (1− *f_p_*)*^x^* ≃ 1 − *x*· *f_p_* to avoid underflow for the computation. In this evaluation, we calculate the probability (6) based on the assumption that both the user and the server create fingerprints by using the same model which generates each bit independently according to the same Bernoulli distribution with probability *w* for the occurrence of the true bit. We notice that this simple assumption is a rather rough approximation to the real problem where bit generation is not always independent and *w* is not equal for all the bits; however, we still think that this evaluation reflects an important tendency of the effect of thresholding and helps us to understand the output privacy of the proposed protocol.

Now, the probability (6) and 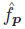 are denoted by *w* as follows.

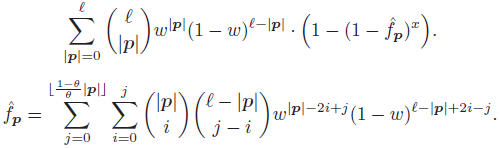

For the calculation of the probability (6) in the Thresholding largely improves database privacy section, we used *w* = 0.28 which is the average ratio for the occurrence of the true bit over all 166 MACCS keys stored in ChEMBL.

### Algorithm 2 Full description of the attack algorithm by using a hit query

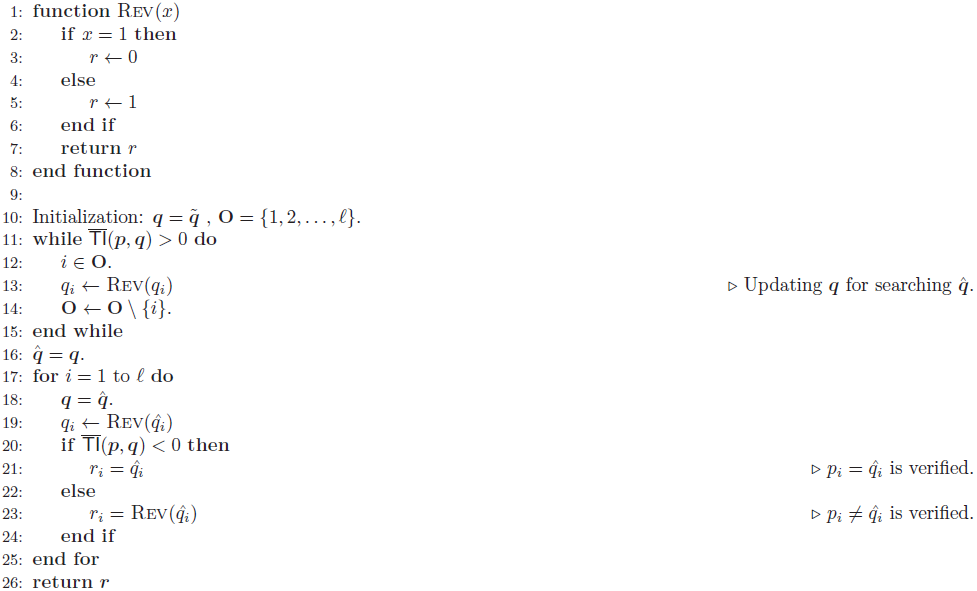

## F Distribution of *w* used in the experiments

In the Security analyses for padding dummies, we tested our method on four different distribution ***w***. Here we list those three distributions. *w_i_* is shown in ascending order by *i*.

- ***w***^ChEMBL−177159^: 0 , 0 , 0 , 0 , 0 , 0 , 0 , 0 , 0 , 0 , 0 , 0 , 0 , 0 , 0 , 0 , 0 , 0 , 0 , 0 , 0 , 0 , 0 , 0 , 0 , 0 , 0 , 0 , 0 , 0 , 0 , 0 , 0 , 0 , 0 , 0 , 0 , 0 , 0 , 0 , 0 , 0 , 0 , 0 , 0 , 0 , 0 , 0 , 0 , 0 , 0 , 0 , 0 , 0 , 0 , 0 , 0 , 0 , 0 , 0 , 0 , 0 , 0 , 0 , 0 , 0 , 0 , 0 , 0 , 0 , 0 , 0 , 0 , 0 , 0 , 0 , 0 , 0 , 0 , 0 , 0 , 0 , 0 , 0 , 0 , 0 , 0 , 0 , 0 , 0 , 0 , 0 , 0 , 0 , 0 , 0 , 0 , 0 , 0 , 0 , 0 , 0 , 0 , 0 , 0 , 0 , 0 , 0 , 0 , 0 , 0 , 0 , 0 , 0 , 0 , 0 , 0 , 0 , 0 , 0 , 0 , 0 , 0 , 0 , 0 , 0 , 0 , 0 , 0 , 0 , 0 , 0 , 0 , 0 , 0 , 0 , 0 , 0 , 0 , 0 , 0 , 0 , 0 , 0 , 0 , 0 , 0 , 0 , 0 , 0 , 0 , 0 , 0 , 0 , 0 , 0 , 0 , 0 , 0 , 0 , 0 , 0 , 0 , 0 , 0 , 0 , 0 , 0 , 0 , 0 , 0 , 0 , 0 , 0 , 0 , 0 , 0 , 0 , 0 , 0 , 0 , 0 , 0 , 0 , 0 , 0 , 0 , 0 , 0 , 0 , 0 , 0 , 0 , 0 , 0 , 0 , 0 , 0 , 0 , 0 , 0 , 0 , 0 , 0 , 0 , 0 , 0 , 0 , 0 , 0 , 0 , 0 , 0 , 0 , 0 , 0 , 0 , 0 , 0 , 0 , 0 , 0 , 0 , 0 , 0 , 0 , 0 , 0 , 0 , 0 , 0 , 0 , 0 , 0 , 0 , 0 , 0 , 0 , 0 , 0 , 0 , 0 , 0 , 0 , 0 , 0 , 0 , 0 , 0 , 0 , 0 , 0 , 0 , 0 , 0 , 0 , 0 , 0 , 0 , 0 , 0 , 0 , 0 , 0 , 0 , 0 , 0 , 0 , 0 , 0 , 0 , 0 , 0 , 0 , 0 , 0 , 0 , 0 , 0 , 0 , 0 , 0 , 0 , 0 , 0 , 0 , 0 , 0 , 0 , 0 , 0 , 0 , 0 , 0 , 0 , 0 , 0 , 0 , 0 , 0 , 0 , 0 , 0 , 0 , 0 , 0 , 0 , 0 , 0 , 0 , 0 , 0 , 0 , 0 , 0 , 1.54758e-06 , 0 , 0 , 0 , 7.73788e-07 , 0 , 0 , 0 , 0 , 0 , 0 , 0 , 7.73788e-07 , 7.73788e-07 , 7.73788e-07 , 0 , 0 , 0 , 0 , 7.73788e-07 , 7.73788e-07 , 3.09515e-06 , 0 , 1.54758e-06 , 7.73788e-07 , 2.32136e-06 , 1.54758e-06 , 3.09515e-06 , 3.09515e-06 , 3.86894e-06 , 4.64273e-06 , 3.09515e-06 , 3.86894e-06 , 3.86894e-06 , 4.64273e-06 , 3.09515e-06 , 6.96409e-06 , 1.31544e-05 , 8.51167e-06 , 9.28545e-06 , 7.73788e-06 , 1.77971e-05 , 2.24398e-05 , 1.31544e-05 , 2.08923e-05 , 2.47612e-05 , 3.40467e-05 , 3.6368e-05 , 2.70826e-05 , 4.33321e-05 , 5.41651e-05 , 4.64273e-05 , 5.49389e-05 , 7.50574e-05 , 7.27361e-05 , 8.51167e-05 , 8.43429e-05 , 0.000119937 , 0.000104461 , 0.000126127 , 0.000141603 , 0.000186483 , 0.000153984 , 0.000182614 , 0.000177197 , 0.000227494 , 0.00022672 , 0.000256898 , 0.000267731 , 0.000293266 , 0.000297908 , 0.000321896 , 0.000332729 , 0.000372966 , 0.000404691 , 0.000425583 , 0.000419393 , 0.000518438 , 0.000550937 , 0.00058421 , 0.000577246 , 0.000625221 , 0.000729682 , 0.00071498 , 0.000780752 , 0.00081712 , 0.000869737 , 0.000937057 , 0.00101211 , 0.0010067 , 0.00109181 , 0.00114907 , 0.00128604 , 0.0011986 , 0.00134175 , 0.00147097 , 0.00151585 , 0.00150811 , 0.00168221 , 0.00178049 , 0.00186947 , 0.00185864 , 0.00201185 , 0.00220916 , 0.00213333 , 0.00223779 , 0.0024916 , 0.00250398 , 0.00265719 , 0.00266493 , 0.00291254 , 0.00304485 , 0.00307968 , 0.00322747 , 0.00337294 , 0.00349288 , 0.00360353 , 0.00366156 , 0.00376448 , 0.00405155 , 0.00412506 , 0.00421482 , 0.00449958 , 0.00457309 , 0.00466826 , 0.00485784 , 0.00492206 , 0.00512712 , 0.00515033 , 0.00539175 , 0.00550705 , 0.00555967 , 0.00578019 , 0.00594888 , 0.00602935 , 0.00623209 , 0.00642399 , 0.00639381 , 0.00655553 , 0.00680778 , 0.00679386 , 0.00693082 , 0.0071645 , 0.0071792 , 0.00748717 , 0.00753282 , 0.00765276 , 0.00783847 , 0.00799168 , 0.00798936 , 0.00861303 , 0.00833292 , 0.00837703 , 0.00852327 , 0.00871672 , 0.00876624 , 0.00876083 , 0.00887225 , 0.00919105 , 0.00921194 , 0.00931021 , 0.0092723 , 0.00919337 , 0.00936515 , 0.00940307 , 0.00943093 , 0.00955318 , 0.00959961 , 0.00945337 , 0.00970562 , 0.00951295 , 0.00945259 , 0.0094642 , 0.00980776 , 0.009653 , 0.00967699 , 0.00978068 , 0.00961199 , 0.00954312 , 0.00929551 , 0.00932646 , 0.00939765 , 0.00922432 , 0.00920266 , 0.0093845 , 0.0089032 , 0.00894653 , 0.00881035 , 0.00881112 , 0.00884053 , 0.00853643 , 0.00861226 , 0.00863238 , 0.00822923 , 0.00802341 , 0.00798626 , 0.00791508 , 0.00782609 , 0.0074833 , 0.0074864 , 0.00742527 , 0.00736491 , 0.00707629 , 0.00674433 , 0.00688284 , 0.00659809 , 0.00653077 , 0.00627465 , 0.00622435 , 0.00611215 , 0.00592644 , 0.00589704 , 0.0057268 , 0.00568425 , 0.00537396 , 0.00527955 , 0.00541032 , 0.005107 , 0.00486635 , 0.00487254 , 0.00474332 , 0.00466052 , 0.00445392 , 0.00441291 , 0.00432238 , 0.00431541 , 0.00425119 , 0.00411346 , 0.00398888 , 0.00406084 , 0.00392465 , 0.00376525 , 0.00395173 , 0.00368942 , 0.00352925 , 0.00362674 , 0.00342014 , 0.00345187 , 0.00335901 , 0.00310908 , 0.00307581 , 0.00295587 , 0.00299224 , 0.0028398 , 0.00283129 , 0.00269742 , 0.00272373 , 0.00253957 , 0.00258522 , 0.00242196 , 0.0023763 , 0.00234845 , 0.00238481 , 0.00212327 , 0.00216428 , 0.00196619 , 0.00224089 , 0.0020103 , 0.00187876 , 0.00172168 , 0.00182614 , 0.00163037 , 0.0016265 , 0.00153674 , 0.00144698 , 0.00133711 , 0.00132163 , 0.00127752 , 0.00124812 , 0.00101985 , 0.0011336 , 0.00109336 , 0.0010214 , 0.000857357 , 0.000885213 , 0.000829501 , 0.000877475 , 0.000793132 , 0.000735872 , 0.000639922 , 0.000667005 , 0.000578019 , 0.000528497 , 0.000483617 , 0.000435643 , 0.000486713 , 0.000421714 , 0.00036368 , 0.000362906 , 0.000361359 , 0.000272373 , 0.000304099 , 0.000225946 , 0.000290944 , 0.000232136 , 0.000201959 , 0.000188804 , 0.000215887 , 0.000181066 , 0.000158626 , 0.00015321 , 0.000153984 , 0.000125354 , 0.000110652 , 0.000106009 , 8.51167e-05 , 0.000102914 , 9.44021e-05 , 7.6605e-05 , 6.26768e-05 , 7.73788e-05 , 5.88079e-05 , 4.48797e-05 , 5.49389e-05 , 5.64865e-05 , 4.10108e-05 , 2.01185e-05 , 3.79156e-05 , 2.47612e-05 , 2.5535e-05 , 2.01185e-05 , 3.40467e-05 , 1.62495e-05 , 1.62495e-05 , 1.85709e-05 , 6.1903e-06 , 1.0833e-05 , 3.86894e-06 , 4.64273e-06 , 3.09515e-06 , 7.73788e-06 , 2.32136e-06 , 3.86894e-06 , 6.96409e-06 , 1.54758e-06 , 4.64273e-06 , 7.73788e-07 , 0 , 0 , 7.73788e-07 , 3.86894e-06 , 7.73788e-07 , 1.54758e-06 , 0 , 7.73788e-07 , 2.32136e-06 , 7.73788e-07 , 2.32136e-06 , 0 , 2.32136e-06 , 0 , 7.73788e-07 , 0 , 0 , 0 , 0 , 0 , 0 , 0 , 0 , 0 , 0 , 0 , 0 , 0 , 7.73788e-07 , 0 , 0 , 0 , 0 , 0 , 0 , 0 , 0 , 0 , 0 , 0 , 0 , 0 , 0 , 0 , 0 , 0 , 7.73788e-07 , 0 , 0 , 0 , 0 , 0 , 0 , 0 , 0 , 0 , 0 , 0 , 0 , 0 , 0 , 0 , 0 , 0 , 0 , 0 , 0 , 0 , 0 , 0 , 0 , 0 , 0 , 0 , 0 , 0 , 0 , 0 , 0 , 0 , 0 , 0 , 0 , 0 , 0 , 0 , 0 , 0 , 0 , 0 , 0 , 0 , 0 , 0 , 0 , 0 , 0 , 0 , 0 , 0 , 0 , 0 , 0 , 0 , 0 , 0 , 0 , 0 , 0 , 0 , 0 , 0 , 0 , 0 , 0 , 0 , 0 , 0 , 0 , 0 , 0 , 0 , 0 , 0 , 0 , 0 , 0 , 0 , 0 , 0 , 0 , 0 , 0 , 0 , 0 , 0 , 0 , 0 , 0 , 0 , 0 , 0 , 0 , 0 , 0 , 0 , 0 , 0 , 0 , 0 , 0 , 0 , 0 , 0 , 0 , 0 , 0 , 0 , 0 , 0 , 0 , 0 , 0 , 0 , 0 , 0 , 0 , 0 , 0 , 0 , 0 , 0 , 0 , 0 , 0 , 0 , 0 , 0 , 0
- ***w***^ChEMBL−265935^ : 0 , 0 , 0 , 0 , 0 , 0 , 0 , 0 , 0 , 0 , 0 , 0 , 0 , 0 , 0 , 0 , 0 , 0 , 0 , 0 , 0 , 0 , 0 , 0 , 0 , 0 , 0 , 0 , 0 , 0 , 0 , 0 , 0 , 0 , 0 , 0 , 0 , 0 , 0 , 0 , 0 , 0 , 0 , 0 , 0 , 0 , 0 , 0 , 0 , 0 , 0 , 0 , 0 , 0 , 0 , 0 , 0 , 0 , 0 , 0 , 0 , 0 , 0 , 0 , 0 , 0 , 0 , 0 , 0 , 0 , 0 , 0 , 0 , 0 , 0 , 0 , 0 , 0 , 0 , 0 , 0 , 0 , 0 , 0 , 0 , 0 , 0 , 0 , 0 , 0 , 0 , 0 , 0 , 0 , 0 , 0 , 0 , 0 , 0 , 0 , 0 , 0 , 0 , 0 , 0 , 0 , 0 , 0 , 0 , 0 , 0 , 0 , 0 , 0 , 0 , 0 , 0 , 0 , 0 , 0 , 0 , 0 , 0 , 0 , 0 , 0 , 0 , 0 , 0 , 0 , 0 , 0 , 0 , 0 , 0 , 0 , 0 , 0 , 0 , 0 , 0 , 0 , 0 , 0 , 0 , 0 , 0 , 0 , 0 , 0 , 0 , 0 , 0 , 0 , 0 , 0 , 0 , 0 , 0 , 0 , 0 , 0 , 0 , 0 , 0 , 0 , 0 , 0 , 0 , 0 , 0 , 0 , 0 , 0 , 0 , 0 , 0 , 0 , 0 , 0 , 0 , 0 , 0 , 0 , 0 , 0 , 0 , 0 , 0 , 0 , 0 , 0 , 0 , 0 , 0 , 0 , 0 , 0 , 0 , 0 , 0 , 0 , 0 , 0 , 0 , 0 , 0 , 0 , 0 , 0 , 0 , 0 , 0 , 0 , 0 , 0 , 0 , 0 , 0 , 0 , 0 , 0 , 0 , 0 , 0 , 0 , 0 , 0 , 0 , 0 , 0 , 0 , 0 , 0 , 0 , 0 , 0 , 0 , 0 , 0 , 0 , 0 , 0 , 0 , 0 , 0 , 0 , 0 , 0 , 0 , 0 , 0 , 0 , 0 , 0 , 0 , 0 , 0 , 0 , 0 , 0 , 0 , 0 , 0 , 0 , 0 , 0 , 0 , 0 , 0 , 0 , 0 , 0 , 0 , 0 , 0 , 0 , 0 , 0 , 0 , 0 , 0 , 0 , 0 , 0 , 0 , 0 , 0 , 0 , 0 , 0 , 0 , 0 , 0 , 0 , 0 , 0 , 0 , 0 , 0 , 0 , 0 , 0 , 0 , 0 , 0 , 0 , 0 , 0 , 0 , 0 , 0 , 0 , 0 , 0 , 1.54758e-06 , 0 , 0 , 0 , 7.73788e-07 , 0 , 0 , 0 , 0 , 0 , 0 , 0 , 7.73788e-07 , 7.73788e-07 , 7.73788e-07 , 0 , 0 , 0 , 0 , 7.73788e-07 , 7.73788e-07 , 3.09515e-06 , 0 , 1.54758e-06 , 7.73788e-07 , 2.32136e-06 , 1.54758e-06 , 3.09515e-06 , 3.09515e-06 , 3.86894e-06 , 4.64273e-06 , 3.09515e-06 , 3.86894e-06 , 3.86894e-06 , 4.64273e-06 , 3.09515e-06 , 6.96409e-06 , 1.31544e-05 , 8.51167e-06 , 9.28545e-06 , 7.73788e-06 , 1.77971e-05 , 2.24398e-05 , 1.31544e-05 , 2.08923e-05 , 2.47612e-05 , 3.40467e-05 , 3.6368e-05 , 2.70826e-05 , 4.33321e-05 , 5.41651e-05 , 4.64273e-05 , 5.49389e-05 , 7.50574e-05 , 7.27361e-05 , 8.51167e-05 , 8.43429e-05 , 0.000119937 , 0.000104461 , 0.000126127 , 0.000141603 , 0.000186483 , 0.000153984 , 0.000182614 , 0.000177197 , 0.000227494 , 0.00022672 , 0.000256898 , 0.000267731 , 0.000293266 , 0.000297908 , 0.000321896 , 0.000332729 , 0.000372966 , 0.000404691 , 0.000425583 , 0.000419393 , 0.000518438 , 0.000550937 , 0.00058421 , 0.000577246 , 0.000625221 , 0.000729682 , 0.00071498 , 0.000780752 , 0.00081712 , 0.000869737 , 0.000937057 , 0.00101211 , 0.0010067 , 0.00109181 , 0.00114907 , 0.00128604 , 0.0011986 , 0.00134175 , 0.00147097 , 0.00151585 , 0.00150811 , 0.00168221 , 0.00178049 , 0.00186947 , 0.00185864 , 0.00201185 , 0.00220916 , 0.00213333 , 0.00223779 , 0.0024916 , 0.00250398 , 0.00265719 , 0.00266493 , 0.00291254 , 0.00304485 , 0.00307968 , 0.00322747 , 0.00337294 , 0.00349288 , 0.00360353 , 0.00366156 , 0.00376448 , 0.00405155 , 0.00412506 , 0.00421482 , 0.00449958 , 0.00457309 , 0.00466826 , 0.00485784 , 0.00492206 , 0.00512712 , 0.00515033 , 0.00539175 , 0.00550705 , 0.00555967 , 0.00578019 , 0.00594888 , 0.00602935 , 0.00623209 , 0.00642399 , 0.00639381 , 0.00655553 , 0.00680778 , 0.00679386 , 0.00693082 , 0.0071645 , 0.0071792 , 0.00748717 , 0.00753282 , 0.00765276 , 0.00783847 , 0.00799168 , 0.00798936 , 0.00861303 , 0.00833292 , 0.00837703 , 0.00852327 , 0.00871672 , 0.00876624 , 0.00876083 , 0.00887225 , 0.00919105 , 0.00921194 , 0.00931021 , 0.0092723 , 0.00919337 , 0.00936515 , 0.00940307 , 0.00943093 , 0.00955318 , 0.00959961 , 0.00945337 , 0.00970562 , 0.00951295 , 0.00945259 , 0.0094642 , 0.00980776 , 0.009653 , 0.00967699 , 0.00978068 , 0.00961199 , 0.00954312 , 0.00929551 , 0.00932646 , 0.00939765 , 0.00922432 , 0.00920266 , 0.0093845 , 0.0089032 , 0.00894653 , 0.00881035 , 0.00881112 , 0.00884053 , 0.00853643 , 0.00861226 , 0.00863238 , 0.00822923 , 0.00802341 , 0.00798626 , 0.00791508 , 0.00782609 , 0.0074833 , 0.0074864 , 0.00742527 , 0.00736491 , 0.00707629 , 0.00674433 , 0.00688284 , 0.00659809 , 0.00653077 , 0.00627465 , 0.00622435 , 0.00611215 , 0.00592644 , 0.00589704 , 0.0057268 , 0.00568425 , 0.00537396 , 0.00527955 , 0.00541032 , 0.005107 , 0.00486635 , 0.00487254 , 0.00474332 , 0.00466052 , 0.00445392 , 0.00441291 , 0.00432238 , 0.00431541 , 0.00425119 , 0.00411346 , 0.00398888 , 0.00406084 , 0.00392465 , 0.00376525 , 0.00395173 , 0.00368942 , 0.00352925 , 0.00362674 , 0.00342014 , 0.00345187 , 0.00335901 , 0.00310908 , 0.00307581 , 0.00295587 , 0.00299224 , 0.0028398 , 0.00283129 , 0.00269742 , 0.00272373 , 0.00253957 , 0.00258522 , 0.00242196 , 0.0023763 , 0.00234845 , 0.00238481 , 0.00212327 , 0.00216428 , 0.00196619 , 0.00224089 , 0.0020103 , 0.00187876 , 0.00172168 , 0.00182614 , 0.00163037 , 0.0016265 , 0.00153674 , 0.00144698 , 0.00133711 , 0.00132163 , 0.00127752 , 0.00124812 , 0.00101985 , 0.0011336 , 0.00109336 , 0.0010214 , 0.000857357 , 0.000885213 , 0.000829501 , 0.000877475 , 0.000793132 , 0.000735872 , 0.000639922 , 0.000667005 , 0.000578019 , 0.000528497 , 0.000483617 , 0.000435643 , 0.000486713 , 0.000421714 , 0.00036368 , 0.000362906 , 0.000361359 , 0.000272373 , 0.000304099 , 0.000225946 , 0.000290944 , 0.000232136 , 0.000201959 , 0.000188804 , 0.000215887 , 0.000181066 , 0.000158626 , 0.00015321 , 0.000153984 , 0.000125354 , 0.000110652 , 0.000106009 , 8.51167e-05 , 0.000102914 , 9.44021e-05 , 7.6605e-05 , 6.26768e-05 , 7.73788e-05 , 5.88079e-05 , 4.48797e-05 , 5.49389e-05 , 5.64865e-05 , 4.10108e-05 , 2.01185e-05 , 3.79156e-05 , 2.47612e-05 , 2.5535e-05 , 2.01185e-05 , 3.40467e-05 , 1.62495e-05 , 1.62495e-05 , 1.85709e-05 , 6.1903e-06 , 1.0833e-05 , 3.86894e-06 , 4.64273e-06 , 3.09515e-06 , 7.73788e-06 , 2.32136e-06 , 3.86894e-06 , 6.96409e-06 , 1.54758e-06 , 4.64273e-06 , 7.73788e-07 , 0 , 0 , 7.73788e-07 , 3.86894e-06 , 7.73788e-07 , 1.54758e-06 , 0 , 7.73788e-07 , 2.32136e-06 , 7.73788e-07 , 2.32136e-06 , 0 , 2.32136e-06 , 0 , 7.73788e-07 , 0 , 0 , 0 , 0 , 0 , 0 , 0 , 0 , 0 , 0 , 0 , 0 , 0 , 7.73788e-07 , 0 , 0 , 0 , 0 , 0 , 0 , 0 , 0 , 0 , 0 , 0 , 0 , 0 , 0 , 0 , 0 , 0 , 7.73788e-07 , 0 , 0 , 0 , 0 , 0 , 0 , 0 , 0 , 0 , 0 , 0 , 0 , 0 , 0 , 0 , 0 , 0 , 0 , 0 , 0 , 0 , 0 , 0 , 0 , 0 , 0 , 0 , 0 , 0 , 0 , 0 , 0 , 0 , 0 , 0 , 0 , 0 , 0 , 0 , 0 , 0 , 0 , 0 , 0 , 0 , 0 , 0 , 0 , 0 , 0 , 0 , 0 , 0 , 0 , 0 , 0 , 0 , 0 , 0 , 0 , 0 , 0 , 0 , 0 , 0 , 0 , 0 , 0 , 0 , 0 , 0 , 0 , 0 , 0 , 0 , 0 , 0 , 0 , 0 , 0 , 0 , 0 , 0 , 0 , 0 , 0 , 0 , 0 , 0 , 0 , 0 , 0 , 0 , 0 , 0 , 0 , 0 , 0 , 0 , 0 , 0 , 0 , 0 , 0 , 0 , 0 , 0 , 0 , 0 , 0 , 0 , 0 , 0 , 0 , 0 , 0 , 0 , 0 , 0 , 0 , 0 , 0 , 0 , 0 , 0 , 0 , 0 , 0 , 0 , 0 , 0 , 0
- ***w***^rand^: 0.00216606 , 0.00168472 , 0.00120337 , 0.00264741 , 0.00120337 , 0.000722022 , 0.000962696 , 0.00120337 , 0.000481348 , 0.000481348 , 0.000481348 , 0.000481348 , 0.00120337 , 0.00120337 , 0.000962696 , 0.000722022 , 0.000481348 , 0.00144404 , 0.000962696 , 0.000962696 , 0.00168472 , 0.00192539 , 0.000722022 , 0.00144404 , 0.000962696 , 0.000962696 , 0.00120337 , 0.000722022 , 0.000962696 , 0.000722022 , 0.00144404 , 0.000962696 , 0.000962696 , 0.00192539 , 0.00192539 , 0.00144404 , 0.00120337 , 0.000962696 , 0.00144404 , 0.00168472 , 0.000481348 , 0.000962696 , 0.000962696 , 0 , 0.000962696 , 0.000481348 , 0.00144404 , 0.000962696 , 0.00168472 , 0.00192539 , 0.00192539 , 0.00288809 , 0.000962696 , 0.00192539 , 0.00144404 , 0.00120337 , 0.00120337 , 0.000722022 , 0.000962696 , 0.00120337 , 0.00144404 , 0.000962696 , 0.00120337 , 0.000240674 , 0.00120337 , 0.000962696 , 0.00216606 , 0.000722022 , 0.000962696 , 0.00144404 , 0.000722022 , 0.000481348 , 0.00144404 , 0.000722022 , 0.000962696 , 0.00168472 , 0.00192539 , 0.00120337 , 0.00120337 , 0.00192539 , 0.000962696 , 0.000722022 , 0.00120337 , 0.000962696 , 0.000962696 , 0.000722022 , 0.00120337 , 0.000962696 , 0.00120337 , 0.00144404 , 0.00120337 , 0.000962696 , 0.00120337 , 0.000240674 , 0.000962696 , 0.00168472 , 0.00120337 , 0.000481348 , 0.000722022 , 0.00288809 , 0.00144404 , 0.000962696 , 0.000240674 , 0.00168472 , 0.00168472 , 0.00216606 , 0.000722022 , 0.00168472 , 0.00144404 , 0.00144404 , 0.00120337 , 0.00120337 , 0.000722022 , 0.000962696 , 0.000962696 , 0.00240674 , 0.000722022 , 0.000481348 , 0.00144404 , 0.000722022 , 0.000481348 , 0.00168472 , 0.00240674 , 0.000240674 , 0.000962696 , 0.000962696 , 0.00240674 , 0.00192539 , 0.000962696 , 0.00192539 , 0.00120337 , 0.000240674 , 0.000722022 , 0.00168472 , 0.00168472 , 0.000962696 , 0.000722022 , 0.00144404 , 0.00144404 , 0.00144404 , 0.000722022 , 0.00168472 , 0.00120337 , 0.000722022 , 0.00144404 , 0.00168472 , 0.00192539 , 0.000240674 , 0.00120337 , 0.00144404 , 0.00168472 , 0.000962696 , 0.00144404 , 0.000722022 , 0.000962696 , 0.000481348 , 0.00120337 , 0.00120337 , 0.000962696 , 0.000481348 , 0.000722022 , 0.00168472 , 0.000481348 , 0.000962696 , 0.000962696 , 0.000962696 , 0.00120337 , 0.000962696 , 0.00216606 , 0.00144404 , 0.00144404 , 0.000722022 , 0.00192539 , 0.000240674 , 0.000962696 , 0.00168472 , 0.00216606 , 0.000962696 , 0.00168472 , 0.00216606 , 0.000722022 , 0.00120337 , 0.00144404 , 0.000962696 , 0.00144404 , 0.00192539 , 0.000722022 , 0.000481348 , 0.000722022 , 0.00168472 , 0.000722022 , 0.000722022 , 0.000481348 , 0.00120337 , 0.00120337 , 0.00120337 , 0.00192539 , 0.000481348 , 0.00192539 , 0.00120337 , 0.00120337 , 0.000962696 , 0.000481348 , 0.000240674 , 0.000722022 , 0.000722022 , 0.00168472 , 0.000481348 , 0.000240674 , 0.00120337 , 0.00144404 , 0.00168472 , 0.00168472 , 0.00120337 , 0.00120337 , 0.00144404 , 0.000722022 , 0.000962696 , 0.00144404 , 0.00120337 , 0.00168472 , 0.000962696 , 0.000722022 , 0.000722022 , 0.00144404 , 0.000481348 , 0.000962696 , 0.000722022 , 0.00144404 , 0.00120337 , 0.000722022 , 0.000481348 , 0.00168472 , 0.000962696 , 0.00168472 , 0.000240674 , 0.000962696 , 0.00120337 , 0.00192539 , 0.000962696 , 0.000240674 , 0.00120337 , 0.000962696 , 0.00240674 , 0.00120337 , 0.000240674 , 0.00144404 , 0.00120337 , 0.000481348 , 0.00168472 , 0.000481348 , 0.00120337 , 0.00144404 , 0.00120337 , 0.000962696 , 0.000962696 , 0.00120337 , 0.00120337 , 0.000962696 , 0.000722022 , 0.000722022 , 0.000962696 , 0.000240674 , 0.00168472 , 0.00168472 , 0.00192539 , 0.000962696 , 0.000722022 , 0.000722022 , 0.00144404 , 0.000481348 , 0.00168472 , 0.00168472 , 0.00216606 , 0.00120337 , 0.00120337 , 0.00120337 , 0.00168472 , 0.000962696 , 0.000962696 , 0.000962696 , 0.000481348 , 0.000481348 , 0.00144404 , 0.00192539 , 0 , 0.00216606 , 0.00144404 , 0.000962696 , 0.00144404 , 0.00144404 , 0.000481348 , 0.000962696 , 0.000722022 , 0.00240674 , 0.00120337 , 0.00144404 , 0.00120337 , 0.00144404 , 0.000962696 , 0.00120337 , 0.00144404 , 0.00120337 , 0.00168472 , 0.000722022 , 0.00144404 , 0.00168472 , 0.000240674 , 0.000240674 , 0.00120337 , 0.000722022 , 0.00168472 , 0.000722022 , 0.000722022 , 0.00216606 , 0.00240674 , 0.00216606 , 0.00120337 , 0.00120337 , 0.00120337 , 0.00168472 , 0.00144404 , 0.000722022 , 0.000962696 , 0.00216606 , 0.000481348 , 0.00168472 , 0.000962696 , 0.000481348 , 0.00120337 , 0.00216606 , 0.000722022 , 0.00120337 , 0.00192539 , 0.00144404 , 0.000722022 , 0.00120337 , 0.00264741 , 0.00216606 , 0.00120337 , 0.000962696 , 0.00120337 , 0.000481348 , 0.00144404 , 0.00144404 , 0.00168472 , 0.000962696 , 0.00240674 , 0.000962696 , 0.00144404 , 0.000240674 , 0.000481348 , 0.000962696 , 0.000481348 , 0.000481348 , 0.000962696 , 0.000481348 , 0.00144404 , 0.00168472 , 0.000962696 , 0.00144404 , 0.00216606 , 0.00216606 , 0.00192539 , 0.000962696 , 0.000962696 , 0.000722022 , 0.00144404 , 0.00216606 , 0.00120337 , 0.00168472 , 0.00120337 , 0.00192539 , 0.000962696 , 0.00120337 , 0.000481348 , 0.000481348 , 0.000722022 , 0.000962696 , 0.000962696 , 0.00144404 , 0.00168472 , 0.000240674 , 0.000722022 , 0.00120337 , 0.00120337 , 0.00192539 , 0.000722022 , 0.00216606 , 0.00120337 , 0.000962696 , 0.00192539 , 0.00216606 , 0.00144404 , 0.000722022 , 0.00120337 , 0.00120337 , 0.00120337 , 0.000481348 , 0.000722022 , 0.000962696 , 0.000722022 , 0.00192539 , 0.00144404 , 0.00144404 , 0.000722022 , 0.000962696 , 0.00192539 , 0.000722022 , 0.00120337 , 0.00120337 , 0.000481348 , 0.000481348 , 0.000962696 , 0.00168472 , 0.00120337 , 0.000962696 , 0.000481348 , 0.00120337 , 0.00120337 , 0.000481348 , 0.00144404 , 0.00168472 , 0.000722022 , 0.00168472 , 0.00120337 , 0.00120337 , 0.00144404 , 0.000722022 , 0.00216606 , 0.000962696 , 0.00144404 , 0.000240674 , 0.00144404 , 0.00192539 , 0.00168472 , 0.00144404 , 0.000962696 , 0.000962696 , 0.00192539 , 0.00240674 , 0.000722022 , 0.000481348 , 0.000722022 , 0.00192539 , 0.000962696 , 0.00168472 , 0.00120337 , 0.000481348 , 0.00168472 , 0.000722022 , 0.00144404 , 0.00144404 , 0.00144404 , 0.000722022 , 0.000962696 , 0.00120337 , 0.00168472 , 0.000962696 , 0.00144404 , 0.00144404 , 0.00144404 , 0.00192539 , 0.00216606 , 0.00120337 , 0.00120337 , 0.000722022 , 0.00168472 , 0.00168472 , 0.000962696 , 0.00168472 , 0.00144404 , 0.00144404 , 0.00264741 , 0.000481348 , 0.000962696 , 0.00168472 , 0.00192539 , 0.00168472 , 0.00120337 , 0.00120337 , 0.000722022 , 0.00120337 , 0.000962696 , 0.000962696 , 0.000962696 , 0.00120337 , 0.000962696 , 0.000481348 , 0.000962696 , 0.00144404 , 0.00168472 , 0.00144404 , 0.000722022 , 0.00120337 , 0.00168472 , 0.00120337 , 0.000722022 , 0.000481348 , 0.000962696 , 0.00168472 , 0.00168472 , 0.00192539 , 0.000240674 , 0.000962696 , 0.000722022 , 0.00192539 , 0.000962696 , 0.00168472 , 0.00192539 , 0.000481348 , 0.00192539 , 0.00144404 , 0.000962696 , 0.00168472 , 0.00168472 , 0.000962696 , 0.00120337 , 0.00144404 , 0.00144404 , 0.000962696 , 0.00144404 , 0.00144404 , 0.000962696 , 0.00216606 , 0.00120337 , 0.00120337 , 0.000481348 , 0.00144404 , 0.000722022 , 0.000962696 , 0.000962696 , 0.00192539 , 0.000481348 , 0.000481348 , 0.000481348 , 0.000722022 , 0.00120337 , 0.000481348 , 0.00144404 , 0.000722022 , 0.00120337 , 0.000962696 , 0.00120337 , 0.00144404 , 0.00168472 , 0.00168472 , 0.00144404 , 0.00168472 , 0.000722022 , 0.00144404 , 0.000722022 , 0.00120337 , 0.000481348 , 0.00144404 , 0.00120337 , 0.00168472 , 0.000722022 , 0.000722022 , 0.00144404 , 0.00144404 , 0.000962696 , 0.000962696 , 0.00144404 , 0.00144404 , 0.00168472 , 0.00168472 , 0.00168472 , 0.000722022 , 0.000722022 , 0.000962696 , 0.000962696 , 0.000722022 , 0.00120337 , 0.00120337 , 0.000722022 , 0.00144404 , 0.00168472 , 0.00144404 , 0.000481348 , 0.000240674 , 0.00168472 , 0.000722022 , 0.00120337 , 0.000240674 , 0.000722022 , 0.00120337 , 0.00120337 , 0.00192539 , 0.00120337 , 0.000962696 , 0.00120337 , 0.000962696 , 0.000722022 , 0.000481348 , 0.00168472 , 0.00120337 , 0.00192539 , 0.00120337 , 0.000240674 , 0.00144404 , 0.00120337 , 0.00192539 , 0.00144404 , 0.000962696 , 0.00192539 , 0.00120337 , 0.00144404 , 0.00240674 , 0.00192539 , 0.000481348 , 0.00216606 , 0.000962696 , 0.000481348 , 0.00144404 , 0.00192539 , 0.000962696 , 0.00264741 , 0.000722022 , 0.00120337 , 0.00168472 , 0.00168472 , 0.000962696 , 0.00144404 , 0.00144404 , 0.00168472 , 0.00120337 , 0.00144404 , 0.00144404 , 0.00144404 , 0.00168472 , 0.00144404 , 0.00168472 , 0.000962696 , 0.000240674 , 0.000962696 , 0.000240674 , 0.000722022 , 0.00120337 , 0.00120337 , 0.000240674 , 0.000722022 , 0.00168472 , 0.00120337 , 0.000962696 , 0.00144404 , 0.00120337 , 0.000962696 , 0.00192539 , 0.00144404 , 0.00120337 , 0.00144404 , 0.00216606 , 0.00144404 , 0.00144404 , 0.000962696 , 0.000481348 , 0.00120337 , 0.000962696 , 0.000481348 , 0.00240674 , 0.000722022 , 0.00120337 , 0.00144404 , 0.00120337 , 0.000481348 , 0.000240674 , 0.000481348 , 0.000962696 , 0.00216606 , 0.00144404 , 0.000722022 , 0.000481348 , 0.000962696 , 0.00168472 , 0.000962696 , 0.000722022 , 0.000962696 , 0.00240674 , 0.00120337 , 0.00144404 , 0.00192539 , 0.000962696 , 0.00192539 , 0.000722022 , 0.000722022 , 0.000962696 , 0.00120337 , 0.000722022 , 0.000481348 , 0.000962696 , 0.00240674 , 0.00192539 , 0.000722022 , 0.000962696 , 0.00120337 , 0.00168472 , 0.00120337 , 0.00120337 , 0.00144404 , 0.00168472 , 0.000722022 , 0.00144404 , 0.00168472 , 0.00144404 , 0.00120337 , 0.00144404 , 0.00120337 , 0.00168472 , 0.00120337 , 0.00120337 , 0.00240674 , 0.000722022 , 0.000722022 , 0.00168472 , 0.000962696 , 0.00144404 , 0.00120337 , 0.00144404 , 0.000481348 , 0.00192539 , 0.00192539 , 0.000481348 , 0.000722022 , 0.000722022 , 0.00144404 , 0.000962696 , 0.00120337 , 0.000722022 , 0.00120337 , 0.00216606 , 0.00144404 , 0.00168472 , 0.00168472 , 0.00216606 , 0.00120337 , 0.000962696 , 0.000481348 , 0.00144404 , 0.00168472 , 0.00144404 , 0.000240674 , 0.00120337 , 0.00120337 , 0.00120337 , 0.00240674 , 0.00144404 , 0.00144404 , 0.00144404 , 0.00144404 , 0.000962696 , 0.00120337 , 0.00120337 , 0.000722022 , 0.000481348 , 0.00168472 , 0.000722022 , 0.000962696 , 0.000962696 , 0.000962696 , 0.000481348 , 0.00144404 , 0.00120337 , 0.000240674 , 0.00144404 , 0.000962696 , 0.00120337 , 0.00120337 , 0.00120337 , 0.000722022 , 0.000962696 , 0.00120337 , 0.00120337 , 0.00144404 , 0.00120337 , 0.00168472 , 0.00168472 , 0.000962696 , 0.00144404 , 0.00120337 , 0.00192539 , 0.000722022 , 0.000962696 , 0.000962696 , 0.000722022 , 0.00144404 , 0.00192539 , 0.000962696 , 0.000722022 , 0.00144404 , 0.000722022 , 0.00144404 , 0.000962696 , 0.00144404 , 0.000962696 , 0.00120337 , 0.000722022 , 0.000962696 , 0.000481348 , 0.00120337 , 0.00120337 , 0.00288809 , 0.00120337 , 0.000481348 , 0.00144404 , 0.00192539 , 0.00144404 , 0.00168472 , 0.000962696 , 0.000722022 , 0.00120337 , 0.00168472 , 0.00216606 , 0.00120337 , 0.00168472 , 0.00120337 , 0.000962696 , 0.000481348 , 0.00120337 , 0.000481348 , 0.00168472 , 0.00168472 , 0.000481348 , 0.00144404 , 0.00168472 , 0.00216606 , 0.00192539 , 0.00120337 , 0.00192539 , 0.00144404 , 0.00120337

## References

1 Subbaraman, N.: Flawed arithmetic on drug development costs. Nature Biotechnology 29(5), 381–381(2011)

2 Miller, M.a.: Chemical database techniques in drug discovery. Nature Reviews Drug Discovery 1(3), 220–7 (2002)

3 Schooler, J.: Unpublished results hide the decline effect. Nature 470, 437 (2011)

4 Ostrovsky, R., Skeith, W.E. III.: A survey of single-database private information retrieval: techniques and applications. In: Proceedings of the 10th International Conference on Practice and Theory in Public-key Cryptography. PKC’07, pp. 393–411(2007)

5 Goethals, B., Laur, S., Lipmaa, H., Mielikäinen, T.: On private scalar product computation for privacy-preserving data mining. In: Proceedings of the 7th Annual International Conference on Information Security and Cryptology. ICISC 2004, pp. 104–120(2004)

6 Blundo, C., Cristofaro, E.D., Gasti, P.: EsPRESSo: Efficient Privacy-Preserving Evaluation of Sample Set Similarity. In: Proceedings of Data Privacy Management and Autonomous Spontaneous Security: 7th International Workshop, DPM 2009 and 5th International Workshop, SETOP 2012. DMP/SETOP 2012, pp. 89–103 (2012)

7 Murugesan, M., Jiang, W., Clifton, C., Si, L., Vaidya, J.: Efficient privacy-preserving similar document detection. The VLDB Journal 19(4), 457–475 (2010)

8 Yao, A.C.-C.: How to generate and exchange secrets. In: Proceedings of the 27th Annual Symposium on Foundations of Computer Science. SFCS ’86, pp. 162–167 (1986)

9 Gentry, C.: Fully homomorphic encryption using ideal lattices. In: Proceedings of the 41st Annual ACM Symposium on Theory of Computing. STOC ’09, pp. 169–178 (2009)

10 Togan, M., Plesca, C.: Comparison-based computations over fully homomorphic encrypted data. In: Communications (COMM), 2014 10th International Conference On, pp. 1–6 (2014). doi:10.1109/ICComm.2014.6866760

11 Laur, S., Lipmaa, H.: On private similarity search protocols. In: Proceedings of the 9th Nordic Workshop on Secure IT Systems. NordSec 2004, pp. 73–77 (2004)

12 Popa, R.A., Redfield, C.M.S., Zeldovich, N., Balakrishnan, H.: In: Proceedings of the 23rd ACM Symposium on Operating Systems Principles. SOSP 11, pp. 85–100

13 Martin, Y.C., Kofron, J.L., Traphagen, L.M.: Do structurally similar molecules have similar biological activity? Journal of Medicinal Chemistry 45(19), 4350–4358 (2002)

14 Miller, J.L.: Recent developments in focused library design: targeting gene-families. Current Topics in Medicinal Chemistry 6(1), 19–29 (2006)

15 Curty, R., Tang, J.: Someone’s loss might be your gain: A case of negative results publications in science. In: Proceedings of the American Society for Information Science and Technology. ASISTS, vol. 49 (2012)

16 Gaulton, A., Bellis, L.J., Bento, A.P., Chambers, J., Davies, M., Hersey, A., Light, Y., McGlinchey, S., Michalovich, D., Al-Lazikani, B., Overington, J.P.: ChEMBL: a large-scale bioactivity database for drug discovery. Nucleic Acids Research 40(Database issue), 1100–1107 (2012)

17 Goldwasser, S., Micali, S.: Probabilistic encryption. J. Comput. Syst. Sci. 28(2), 270–299 (1984)

18 Paillier, P.: Public-key cryptosystems based on composite degree residuosity classes. In: Proceedings of the 17th International Conference on Theory and Application of Cryptographic Techniques. EUROCRYPT’99, pp. 223–238 (1999)

19 ElGamal, T.: A public key cryptosystem and a signature scheme based on discrete logarithms. IEEE Transactions on Information Theory 31(4), 469–472 (1985)

20 Goldreich, O.: Foundations of Cryptography: Volume 1. Cambridge University Press, ??? (2001)

21 Sakai, Y., Emura, K., Hanaoka, G., Kawai, Y., Omote, K.: Methods for restricting message space in public-key encryption. IEICE Transactions 96-A(6), 1156–1168 (2013)

22 Tversky, A.: Features of similarity. Psychological Review 84(4), 327–352 (1977)

23 Damgård, I., Fitzi, M., Kiltz, E., Nielsen, J.B., Toft, T.: Unconditionally secure constant-rounds multi-party computation for equality, comparison, bits and exponentiation. In: Proceedings of the 3rd Theory of Cryptography Conference. TCC 2006, pp. 285–304 (2006)

24 C++ Library implementing elliptic curve ElGamal crypto system [19]. https://github.com/aistcrypt/Lifted-ElGamal. URL accessed April 13, 2015 (2015)

25 Ben-david, A., Nisan, N., Pinkas, B.: Fairplaymp: A system for secure multi-party computation. In: Proceedings of ACM Conference on Computer and Communications Security. CCS 2008, pp. 17–21 (2008)

26 Pinkas, B., Schneider, T., Smart, N.P., Williams, S.C.: Secure two-party computation is practical. In: Proceedings of the 15th International Conference on the Theory and Application of Cryptology and Information Security. ASIACRYPT 2009, pp. 250–267 (2009)

27 Kolesnikov, V., Schneider, T.: Improved garbled circuit: Free XOR gates and applications. In: Proceedings of the 35th International Colloquium on Automata, Languages and Programming. ICALP 2008, pp. 486–498 (2008)

28 Henecka, W., Kögl, S., Sadeghi, A., Schneider, T., Wehrenberg, I.: TASTY: tool for automating secure two-party computations. In: Proceedings of the 17th ACM Conference on Computer and Communications Security. CCS 2010, pp. 451–462 (2010)

29 Huang, Y., Evans, D., Katz, J., Malka, L.: Faster secure two-party computation using garbled circuits. In: Proceedings of the 20th USENIX Security Symposium. USENIX 2011 (2011)

30 Huang, Y., Shen, C.-H., Evans, D., Katz, J., Shelat, A.: Efficient secure computation with garbled circuits. In: Proceedings of the 7th International Conference on Information Systems Security. ICISS, pp. 28–48 (2011)

31 Kreuter, B., Shelat, A., Shen, C.: Billion-gate secure computation with malicious adversaries. In: Proceedings of the 21th USENIX Security Symposium. USENIX Security 2012, pp. 285–300 (2012)

32 Williams, A.J., Harland, L., Groth, P., Pettifer, S., Chichester, C., Willighagen, E.L., Evelo, C.T., Blomberg, N., Ecker, G., Goble, C., Mons, B.: Open PHACTS: semantic interoperability for drug discovery. Drug Discovery Today 17(21–22), 1188–1198 (2012)

33 Hunter, J., Stephens, S.: Is open innovation the way forward for big pharma? Nature Reviews Drug Discovery 9(2), 87–88 (2010)

34 Fiat, A., Shamir, A.: How to prove yourself: Practical solutions to identification and signature problems. In: Proceedings on Advances in Cryptology. CRYPTO ’86 (1987)

